# Role of Cytosolic, Tyrosine-Insensitive Prephenate Dehydrogenase in *Medicago truncatula*

**DOI:** 10.1101/768317

**Authors:** Craig A. Schenck, Josh Westphal, Dhileepkumar Jayaraman, Kevin Garcia, Jiangqi Wen, Kirankumar S. Mysore, Jean-Michel Ané, Lloyd W. Sumner, Hiroshi A. Maeda

## Abstract

L-Tyrosine (Tyr) is an aromatic amino acid synthesized *de novo* in plants and microbes downstream of the shikimate pathway. In plants, Tyr and a Tyr pathway intermediate, 4-hydroxyphenylpyruvate (HPP), are precursors to numerous specialized metabolites, which are crucial for plant and human health. Tyr is synthesized in the plastids by a TyrA family enzyme, arogenate dehydrogenase (ADH/TyrA_a_), which is feedback inhibited by Tyr. In addition to ADH enzymes, many legumes possess prephenate dehydrogenases (PDH/TyrA_p_), which are insensitive to Tyr and localized to the cytosol. Yet the role of PDH in legumes is currently unknown. This study isolated and characterized *Tnt1*-transposon mutants of *MtPDH1* (*pdh1*) in *Medicago truncatula* to investigate PDH function. The *pdh1* mutants lacked *PDH* transcript, PDH activity, and displayed little aberrant morphological phenotypes under standard growth conditions providing genetic evidence that *MtPDH1* is responsible for the PDH activity detected in *M. truncatula*. Though plant PDH enzymes and activity have been specifically found in legumes, nodule number and nitrogenase activity of *pdh1* mutants were not significantly reduced compared to wild-type (Wt) during symbiosis with nitrogen-fixing bacteria. Although Tyr levels were not significantly different between Wt and mutants under standard conditions, when carbon flux was increased by shikimate precursor feeding, mutants accumulated significantly less Tyr than Wt. These data suggest that MtPDH1 is involved in Tyr biosynthesis when the shikimate pathway is stimulated, and possibly linked to unidentified legume-specific specialized metabolism.

## INTRODUCTION

L-Tyrosine (Tyr) is an aromatic amino acid synthesized *de novo* in plants and microbes, but not animals; thus, humans must acquire Tyr through their diet or by enzymatic conversion of L-phenylalanine (Phe, Fitzpatrick, 1999). In addition to its involvement in protein synthesis, Tyr and a Tyr-pathway intermediate 4-hydroxyphenylpyruvate (HPP) are the precursors to numerous specialized metabolites crucial for plant and animal health (Schenck and Maeda, 2018). Tyr-derived plant specialized metabolites have roles as photosynthetic electron carriers (plastoquinone, Metz et al., 1989), pollinator attractors (betalain pigments, Strack et al., 2003) and in defense (dhurrin and rosmarinic acid, Møller, 2010; Petersen, 2013). In grasses, Tyr also serves as a precursor to lignin, the main structural polymer in plants (Higuchi et al., 1967; Rosler et al., 1997; Barros et al., 2016). Humans have co-opted some of these natural products to serve nutritional and medicinal roles such as some benzylisoquinoline alkaloids, which have antitussive, analgesic, and antimicrobial activities (Barken et al., 2008; Beaudoin and Facchini, 2014; Kries and O’Connor, 2016), and the antioxidant properties of tocopherols (collectively referred to as vitamin E, Bramley et al., 2000).

The aromatic amino acids (AAAs; Tyr, Phe, and tryptophan [Trp]) are synthesized in the plastids from chorismate, the final product of the shikimate pathway (Tzin and Galili, 2010; Maeda and Dudareva, 2012). Chorismate mutase (CM, EC number 5.4.99.5) catalyzes the committed step of Tyr and Phe synthesis — the isomerization of chorismate into prephenate (Goers and Jensen, 1984; Kuroki and Conn, 1989; Eberhard et al., 1996; Mobley et al., 1999). Prephenate is converted to Tyr via two reactions, oxidative decarboxylation catalyzed by a TyrA dehydrogenase enzyme and transamination. These reactions can occur in either order, leading to alternative Tyr pathways (**Fig. 1**, Schenck and Maeda, 2018). In most microbes, a prephenate-specific TyrA dehydrogenase (PDH/TyrA_p_ EC 1.3.1.13, **Fig. 1**) first converts prephenate into HPP followed by transamination to Tyr via Tyr-aminotransferase (Tyr-AT or TyrB, EC2.6.1.5). In plants, these reactions occur in the reverse order with transamination of prephenate to form arogenate by a plastidic prephenate aminotransferase (PPA-AT, EC 2.6.1.78), followed by oxidative decarboxylation catalyzed by a plastidic arogenate-specific TyrA dehydrogenase (ADH/TyrA_a_ EC 1.3.1.78) producing Tyr (**Fig. 1**). The ADH-mediated Tyr pathway appears to be essential for normal growth and development in plants as indicated by the severe phenotype of *Arabidopsis thaliana adh2/tyra2* knockout mutant, which was further exacerbated by transient suppression of the other *ADH1/TyrA1* gene (de Oliveira et al., 2019). PDH and ADH are the key enzymes in their respective pathways, as they compete for substrates that are shared with Phe biosynthesis, and are generally feedback inhibited by Tyr (Rubin and Jensen, 1979; Gaines et al., 1982; Connelly and Conn, 1986; Rippert and Matringe, 2002).

**Figure 1.**
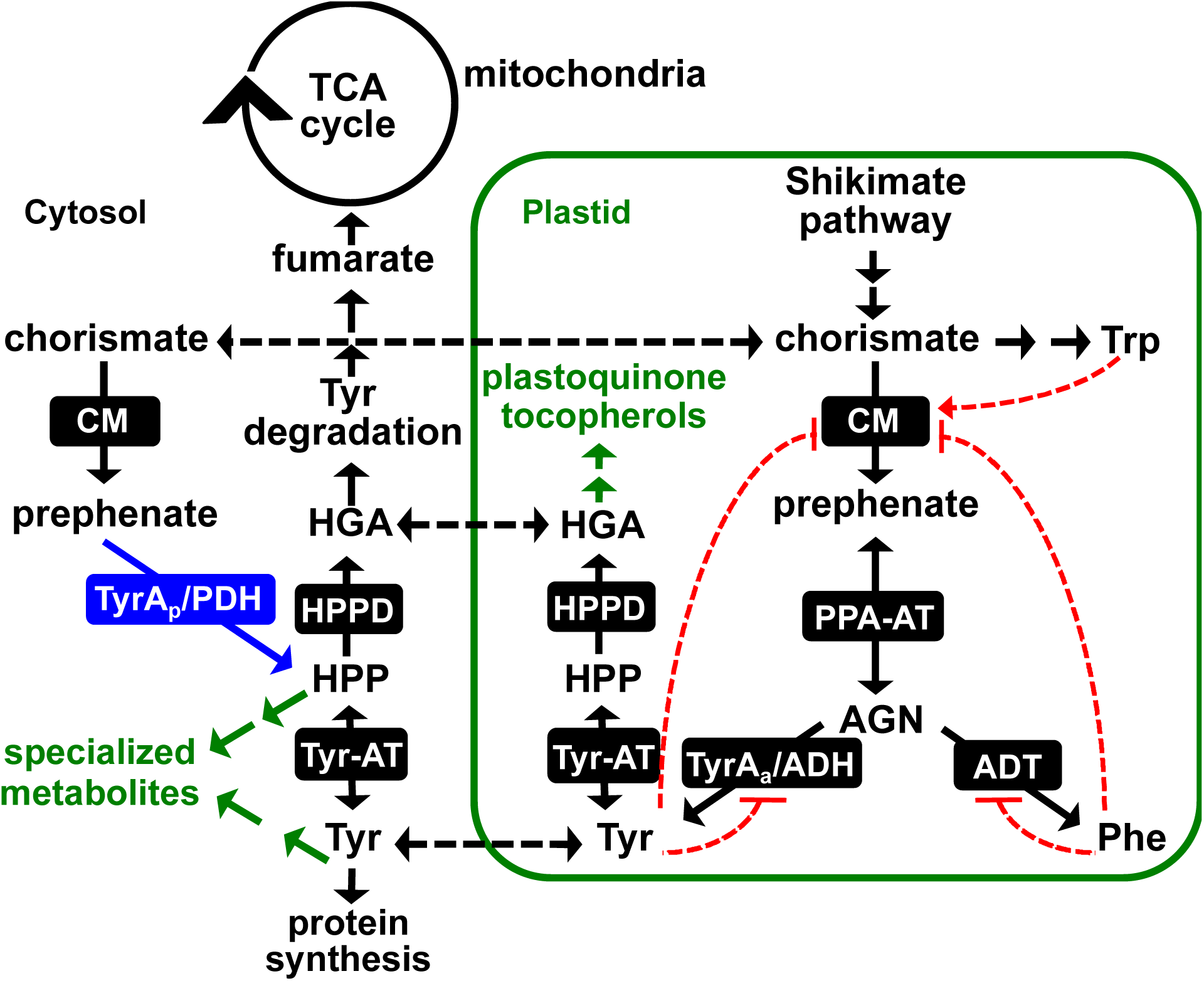
Tyr biosynthesis in legumes. In addition to a highly regulated plastid localized ADH pathway for Tyr biosynthesis, legumes possess a Tyr-insensitive, cytosolic PDH enzyme (blue). Tyr synthesized in the plastids is exported into the cytosol where it can be incorporated into proteins, enter the Tyr degradation pathway to the TCA cycle or serve as a precursor to specialized metabolism. Black dotted lines denote known or potential transport steps, red dotted lines represent feedback regulation with arrows meaning induction and hashes inhibition. AGN, arogenate; ADT, arogenate dehydratase; CM, chorismate mutase; HGA, homogentisate; HPP, 4-hydroxyphenylpyruvate; HPPD, HPP dioxygenase; PPA-AT, prephenate aminotransferase; TyrA_a_/ADH, arogenate dehydrogenase; TyrA_p_/PDH, prephenate dehydrogenase; Tyr-AT, tyrosine aminotransferase.

PDH activity, which is commonly found in microbes, has been detected in tissue extracts of some plants, all restricted to the legume family (Gamborg and Keeley, 1966; Rubin and Jensen, 1979; Siehl, 1999). In soybean, leaf tissue had the highest PDH activity of all analyzed tissues (Schenck et al., 2015). Phylogenetic analyses of plant and microbial TyrAs showed that *PDH* genes are uniquely present in legumes, suggesting that legume PDH enzymes are the result of a recent legume-specific gene duplication event of a plant *ADH* rather than horizontal gene transfer from a *PDH*-possessing rhizobia (Schenck et al., 2015). However, not all legumes possess *PDH*, which suggests gene loss in some legumes (Schenck et al., 2015; Schenck et al., 2017a). PDH recombinant enzymes from *Glycine max* (soybean; GmPDH1 and GmPDH2) and *Medicago truncatula* (Medicago; MtPDH1) preferred prephenate over arogenate as their substrates (Schenck et al., 2015; Schenck et al., 2017a) and, unlike plant ADH enzymes, were insensitive to Tyr inhibition and localized to the cytosol (Schenck et al., 2015; Schenck et al., 2017a). These recently diverged plant ADH and PDH enzymes were used to identify a single amino acid residue (Asn222 of GmPDH1) of TyrA dehydrogenases that switches TyrA substrate specificity and underlies the evolution of legume PDH enzymes (Schenck et al., 2017a; Schenck et al., 2017b). While these biochemical and evolutionary studies established that some legumes have an alternative Tyr-insensitive cytosolic PDH enzyme, its *in vivo* function is unknown.

To address this issue, we hypothesized and tested four non-mutually exclusive *in planta* functions of the PDH enzyme in legumes. **Hypothesis I: PDH functions as a part of a redundant Tyr biosynthetic pathway in the cytosol** (**Fig. 1**). Although AAA biosynthesis is localized to the plastids (Bickel et al., 1978; Jung et al., 1986), some plants including legumes identified cytosolic isoforms of CM and Tyr-AT, which catalyze immediately up- and down-stream steps of PDH and complete the cytosolic Tyr biosynthetic pathway from chorismate (**Fig. 1**) (D’Amato et al., 1984; Eberhard et al., 1996; Ding et al., 2007; Schenck et al., 2015; Wang et al., 2016). Also, a cytosolic Phe pathway was recently identified in plants (Yoo et al., 2013; Qian et al., 2019). **Hypothesis II: PDH is involved in the production of Tyr or HPP-derived metabolite(s) (Fig. 1**). Tyr is the precursor of many plant specialized metabolites (Schenck and Maeda, 2018) and duplicated primary metabolic enzymes can be co-opted to efficiently provide Tyr and HPP precursors to support their downstream specialized metabolism (Weng et al., 2012; Moghe and Last, 2015; Maeda, 2019). **Hypothesis III: PDH is involved in the Tyr catabolism pathway to the tricarboxylic acid (TCA) cycle**. Tyr catabolism proceeds through HPP, the product of PDH, and feeds intermediates (e.g. fumarate) of the TCA cycle (**Fig. 1**). Thus, HPP produced from PDH may be directly incorporated into the Tyr degradation pathway. **Hypothesis IV: PDH is involved in the legume-rhizobia symbiosis.** Many legumes form a symbiotic relationship with rhizobia, and PDH activity is uniquely present in legumes. Furthermore, *MtPDH1* expression is upregulated in the nodules after treatment with nitrate (NO_3_^−^, Benedito et al., 2008) or phosphinothricin (PPT, Seabra et al., 2012), both of which stimulate nodule senescence (Streeter and Wong, 1988; Matamoros et al., 1999; Pérez Guerra et al., 2010). To systematically test the above four hypotheses, this study isolated and characterized *pdh* mutants in the model plant *M. truncatula*, which conveniently has a single *PDH* gene compared with some other legumes that have multiple copies (Schenck et al., 2015; Schenck et al., 2017a) using genetics, biochemistry, metabolomics, histochemical staining, microscopy, and gene expression analyses.

## RESULTS

### Isolation of pdh mutants in Medicago truncatula

To investigate the *in vivo* function of PDH in legumes, two independent homozygous alleles of *Tnt1*-transposon mutants of *M. truncatula* (Tadege et al., 2008) were identified for the *MtPDH1* locus (Mt3g071980) (Cheng et al., 2014). The *pdh1-1* and *pdh1-2* mutants carried a transposon insertion in the first and last exons of the *MtPDH1* gene, respectively (**Fig. 2a**). *MtPDH1* was constitutively expressed across many tissues and under various conditions with the highest and lowest expression being detected in aerial tissues and seeds, respectively (**Supplementary Fig. 1**). Both alleles did not produce any *PDH* transcript in the leaves (**Fig. 2b**). The *M. truncatula* genome contains two *ADH* genes, *MtADH* (Mt4g115980) and *MtncADH* (Mt5g083530) (Schenck et al., 2017a), but neither were enhanced in the *pdh1* mutants (**Fig. 2b**). Consistent with their transcript levels, PDH activity was almost completely abolished in both *pdh1* mutants without significant reduction in ADH activity (**Fig. 2c**). After six weeks of growth, *pdh1-1* showed no phenotypic difference from wild-type (Wt; R108). The *pdh1-2* mutant, on the other hand, had a slight bushy phenotype, which could be due to an unknown secondary transposon insertion(s) (**Fig. 2d**); however, multiple attempts of backcrossing were unsuccessful. These data provide genetic evidence that *MtPDH* is responsible for the PDH activity detected in *M. truncatula*. The minor impacts of eliminating PDH activity on overall plant growth in two independent *pdh1* mutants suggests that the PDH enzyme is not essential during standard growth conditions in *M. truncatula*.

**Figure 2.**
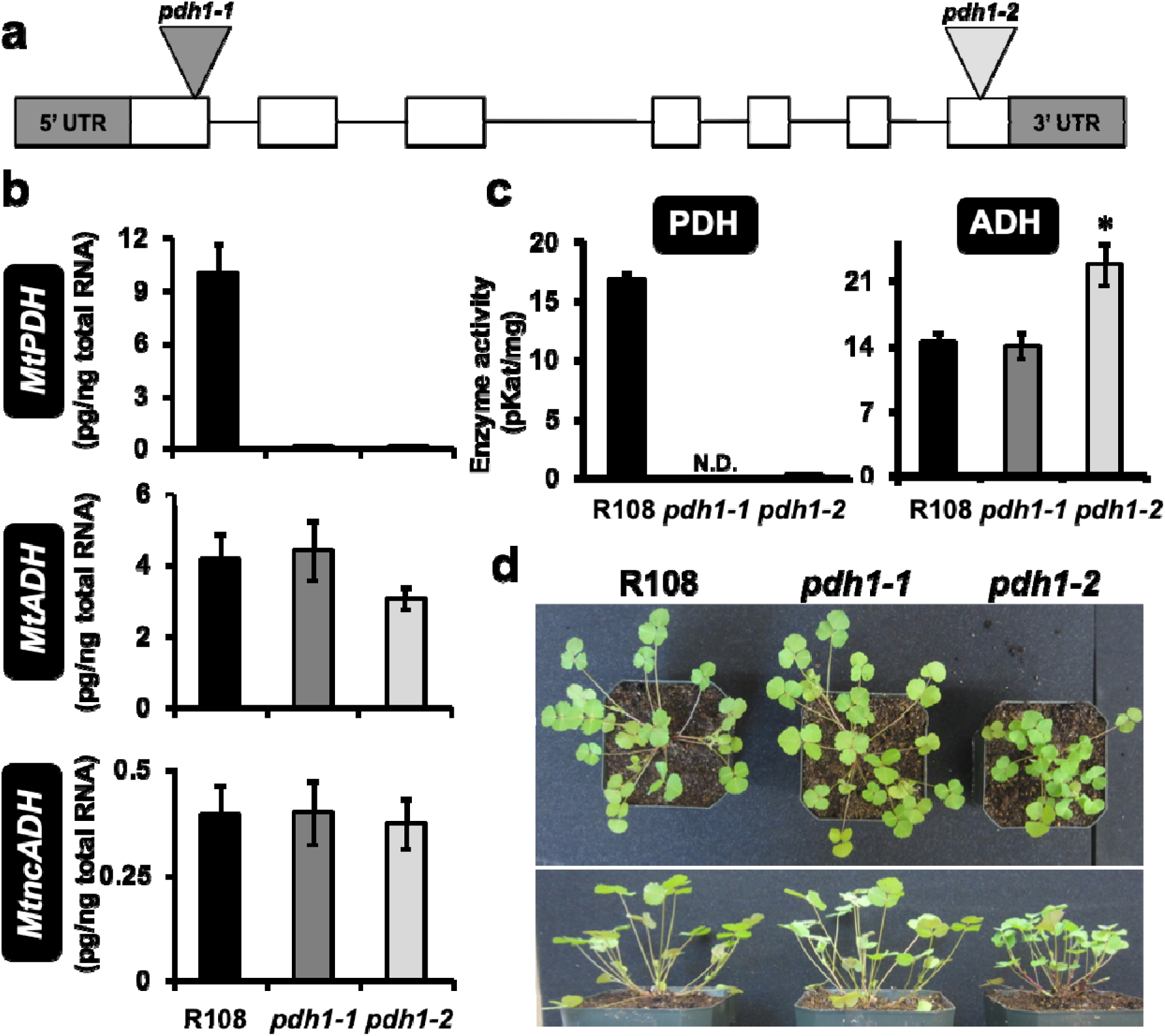
Isolation of *MtPDH1* mutants. **(a)** *MtPDH1* (Mt3g071980) genomic structure, exons (whit boxes) and introns (lines), 5’ and 3’ untranslated regions (UTR, gray boxes). Two *Tnt1*-transposon mutants were isolated with insertions in exon one (*pdh1-1*) and seven (*pdh1-2*). **(b)** *MtPDH1* transcripts were nearly abolished in *pdh1-1* and *pdh1-2*, without effecting expression of either *ADH* homolog. Bars represent average absolute mRNA levels (pg/ng total RNA) ± s.e.m of n = 3 biological replicates. **(c)** PDH and ADH activity from mutants and wild-type (Wt, R108). Bars represent average enzymatic activity (pKat mg) ± s.e.m of n = 3 biological replicates. Significant differences to R108 control are indicated; \**P* ≤ 0.05. **(d)** Phenotype of R108 and mutants after 6-weeks growth under standard conditions.

### Tyr and Tyr-derived compounds are unaltered in pdh mutants during standard growth condition

To test the hypothesis that the PDH pathway serves as a redundant Tyr biosynthetic route (**hypothesis I**), Tyr and Tyr/HPP-derived compounds were analyzed in Wt and mutants. Metabolites were extracted from root and leaf tissue of 6-week-old plants grown under standard conditions. Surprisingly, the levels of Tyr were not significantly different between Wt and mutants in either leaf or root tissue (**Fig. 3a**). Additionally, the levels of Phe and Trp in the leaves and roots were not significantly reduced in mutants as compared to Wt, though a slightly higher Trp level was observed in *pdh1-2* (**Fig. 3a**). Potential effects on tocopherols, HPP-derived metabolites, were also tested (**hypothesis II**), but their levels were not significantly different between Wt and mutants in both leaf and root tissue (**Fig. 3b**). We also performed non-targeted analysis using GC-MS for both polar and non-polar metabolites; however, no consistent differences were observed between Wt and mutants (**Supplementary Table 1**). Together, these data suggest that under standard growth conditions, the lack of the PDH enzyme had no substantial effects on the overall accumulation of aromatic amino acids (AAAs) or Tyr/HPP-derived metabolites analyzed.

**Figure 3.**
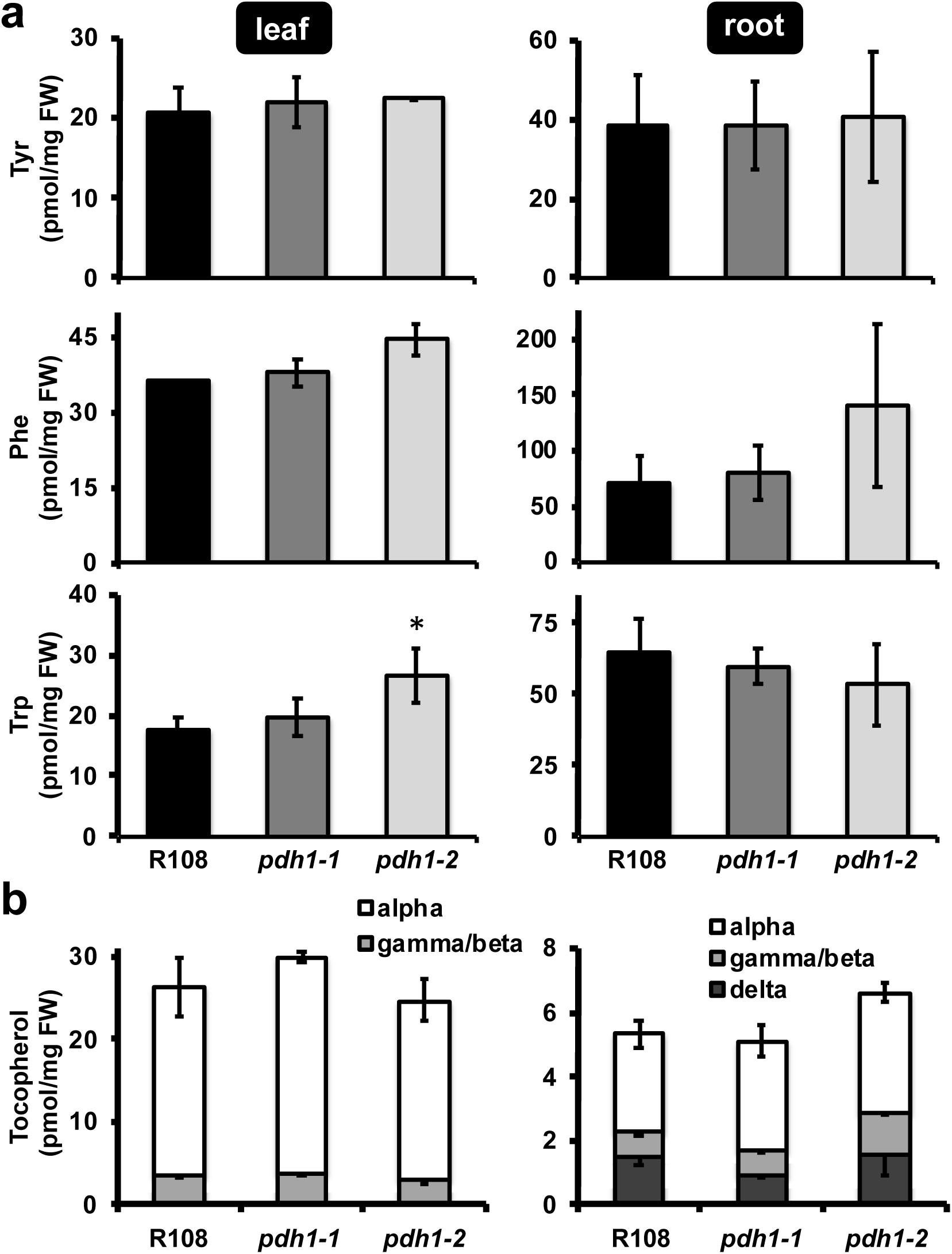
Targeted metabolite analysis of *mtpdh1* mutants and Wt. Leaf and root tissue from 6-week-old plants, grown under standard conditions were used for metabolite extraction. Bars represent average absolute metabolite levels (pmol/mg fresh weight (FW)) ± s.e.m of n = 3 biological replicates. **(a)** Aromatic amino acid levels in leaf and root tissue. Significant differences to Wt (R108) control are indicated; \**P* ≤ 0.05. **(b)** Tocopherol composition and content from leaf and root tissue.

In grasses, upwards of 50% of the total lignin is derived from Tyr, via a Tyr ammonia-lyase (TAL) enzyme (Barros et al., 2016). Since TAL activity has also been detected in some non-grass species, including legumes (Giebel, 1973; Beaudoin-Eagan and Thorpe, 1985; Khan et al., 2003), the legume PDH enzyme may synthesize Tyr that is directly incorporated into the phenylpropanoid pathway for downstream products such as lignin (**hypothesis II**). To test this possibility, stem cross-sections from Wt and mutants were stained using two different techniques, Mäule and phloroglucinol, which can detect potential differences in composition or linkages of lignin (Mitra and Loqué, 2014, Pomar et al., 2002). Neither staining method showed any obvious differences in stem lignification between Wt and mutants (**Supplementary Fig. 2a,b**). When phenylpropanoid intermediates, *p*-coumarate and ferulate, involved in lignin biosynthesis were analyzed by GCMS, they were somewhat reduced in *pdh1-2* but not consistently in *pdh1-1* (**Supplementary Fig. 2c**). Thus, the lack of PDH does not have substantial impacts on lignin biosynthesis in *M. truncatula*.

### Less tocopherols are accumulated in pdh1 than Wt under high light treatment

Under various biotic and abiotic stresses, the shikimate pathway is induced, which often leads to accumulation of AAAs (Dyer et al., 1989; Gilbert et al., 1998; Zhao et al., 1998; Betz et al., 2009). To test the potential role of PDH in Tyr biosynthesis under stress (**hypothesis I**), Wt and *pdh1-1* were subjected to 48-hour high light treatment (Gonzali et al., 2009), which is known to induce production of AAA-derived antioxidants (Collakova and DellaPenna, 2003). The levels of Tyr were not altered, except at 24 hours when Tyr increased slightly in *pdh1-1* compared with Wt (**Supplementary Fig. 3a**). As expected, high light treatment enhanced tocopherol accumulation, but to a significantly lesser extent in *pdh1-1* compared with Wt at both 24 and 48 hours (**Supplementary Fig. 3b**). The levels of anthocyanins, which are also induced under various stresses including high light stress (Collakova and DellaPenna, 2003), were increased after 24 and 48 hours of high light treatment but was not significantly different between Wt and *pdh1-1* (**Supplementary Fig. 3c**). These results show that the lack of PDH negatively impacts the accumulation of HPP-derived tocopherols when their production is induced under high light conditions.

### MtPDH1 is co-expressed with senescence-related genes but pdh1 deficiency has no major impacts on dark-induced senescence

To identify potential processes and pathways that are coordinately regulated with *PDH*, a gene co-expression analysis with *MtPDH1* was performed using the Medicago gene expression atlas (He et al., 2009). *MtPDH1* was co-expressed with genes mainly involved in senescence-related processes (e.g., nucleases, proteases, and lipases **Supplementary Fig. 1**) and the gene encoding HPP dioxygenase (HPPD, (Siehl et al., 2014), a senescence-activated enzyme involved in Tyr catabolism (**Supplementary Fig. 5a**)(Wang et al., 2019). To experimentally test the potential involvement of PDH in senescence, *PDH* gene expression and enzymatic activity were monitored at different developmental stages during natural leaf senescence (**Supplementary Fig. 4a**). Expression of a senescence marker gene (*MtVPE*, Pérez Guerra et al., 2010) was monitored, together with loss of chlorophyll in leaves collected at various developmental times from a single plant, to define stages of leaf senescence (**Supplementary Fig. 4b**). *MtVPE* was basally expressed in fully green leaves (defined here as the S1 stage), and gradually induced upon senescence (defined here as S2, S3, and S4; **Supplementary Fig. 4b**), mirroring the loss of chlorophyll. Fully senescent leaves (S5) were collected but did not yield high-quality RNA and proteins and were unable to be further analyzed. The highest *MtPDH1* expression was detected in green (S1) leaves, and PDH enzymatic activity was not induced upon senescence (**Supplementary Fig. 4c,d**). Thus, *MtPDH1* is not upregulated during natural senescence in the leaves. Similar to *PDH*, expression of the two *ADH* genes in *M. truncatula* did not follow the developmental pattern of leaf senescence (**Supplementary Fig. 4c**). Unlike PDH activity, however, ADH enzymatic activity was gradually induced upon senescence (**Supplementary Fig. 4d**).

Since *MtPDH1* was co-expressed with *MtHPPD* (**Supplementary Fig. 1**), and PDH together with HPPD provides a direct route for catabolism of Tyr to homogentisate, eventually leading to fumarate (**Fig. 1, Supplementary Fig. 5a**), we investigated potential impacts of *pdh1* deficiency in Tyr catabolism (**hypothesis III**). The expression of genes encoding all enzymes of the Tyr catabolic pathway were measured in the mutants and Wt under standard growth conditions (**Supplementary Fig. 5a**, Dixon and Edwards, 2006). The expression of genes encoding the first two steps of the pathway, *HPPD* and homogentisate dioxygenase (*HGO*), were not significantly altered in the mutants compared with Wt. The subsequent step, maleylacetoacetate isomerase (*MAAI*), was induced by 2- and 2.5-fold in *pdh1-1* and *pdh1-2*, respectively (**Supplementary Fig. 5b**), though the final step in the pathway, fumarylacetoacetate hydrolase (*FAH*), showed opposite expression patterns in the two mutants (**Supplementary Fig. 5b**). Thus, no consistent changes in the expression of the Tyr degradation pathway genes, beyond *MAAI* were observed in *pdh1* mutants.

To further examine Tyr catabolism during leaf senescence under artificial, but controlled, conditions excised leaves from Wt and mutants were exposed to an extended dark treatment (Xing and Last, 2017; Wang et al., 2019). Over 7 days, leaves from Wt and mutants responded to dark-induced senescence similarly with no apparent growth phenotypes (**Supplementary Fig. 6a**). Furthermore, there were no significant differences in α-tocopherol levels between Wt and mutants at any time point analyzed (**Supplementary Fig. 6b**). These data together suggest that the lack of PDH had no substantial effects on Tyr catabolism or metabolism under standard growth conditions and at least under the senescence conditions tested here (**Supplementary Figs. 5 & 6**).

### Less Tyr is accumulated in pdh mutants following shikimate feeding

Although the steady-state levels of Tyr were not altered in the mutants (**Fig. 3a**), some of the above data suggest that the PDH enzyme may contribute to Tyr production (**hypothesis I**) at least under some stress conditions (**Supplementary Figs. 1 and 3b**). Thus, we hypothesized that the potential role of PDH in Tyr biosynthesis might be manifested when the carbon flux through the Tyr pathway is elevated. To test this possibility, an intermediate of the shikimate pathway, shikimate, was exogenously fed to Wt and mutants and the levels of AAAs were analysed. Excised leaves were floated in a solution containing shikimate or H_2_O for up to 8 hours, rinsed to remove metabolites in the feeding solution, and leaf metabolites were extracted and analyzed using HPLC and GC-MS. As more prolonged feeding with shikimate led to abnormal leaf phenotypes, 8 hours was selected as an optimal time for increased carbon flux without pleiotropic effects. After 8 hours of feeding, shikimate and Phe levels were increased drastically in all genotypes, suggesting that shikimate was taken up and metabolized by the leaves (**Fig. 4**). Tocopherol levels were generally unaffected upon shikimate feeding, likely because the feeding time was not long enough to convert shikimate into tocopherols (**Fig. 4**). Trp levels were reduced in the mutants compared with Wt after feeding with water, and were induced upon shikimate feeding in one mutant (**Fig. 4**). The unexpected differences observed with Trp levels under standard growth conditions (**Fig. 3**) and the feeding experiments (**Fig. 4**) may reflect unknown stress responses during the feeding experiments. Nevertheless, after 8 hours of feeding, Tyr levels increased by >39-fold in Wt, but only by 16-and 13-fold in *pdh1-1* and *pdh1-2*, respectively (**Fig. 4**). Repeated 8 hour shikimate feeding experiments with *pdh1* mutants and Wt yielded similar Tyr accumulation patterns. The difference in the Tyr accumulation suggests that PDH contributes to Tyr biosynthesis when carbon flow through the shikimate pathway is enhanced.

**Figure 4.**
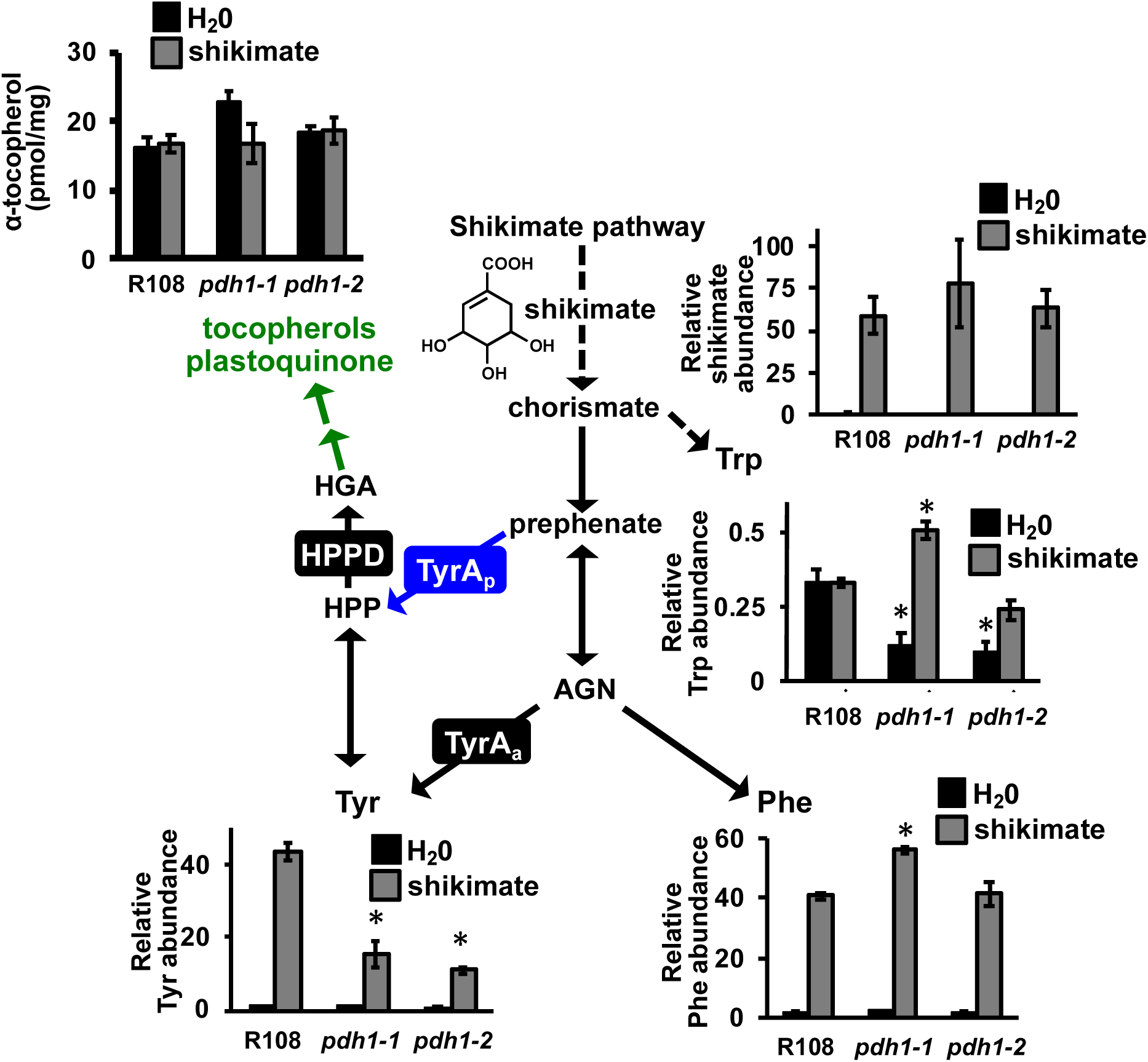
Metabolite analysis after shikimate feeding in Wt and *mtpdh1* mutants. Excised leaves from 6-week-old plants were floated on a solution containing H_2_0 (black bars) or 25 mM shikimate (gray bars) for 8 hours under constant light. Leaves were then used for metabolite analysis using GC-MS (Tyr, Trp, Phe and shikimate) or HPLC (α-tocopherol). Bars represent average relative metabolite abundance of Tyr, Phe, Trp and shikimate ± s.e.m of n = 3 biological replicates. α-tocopherol is shown as the average absolute metabolite abundance in pmol/mg FW ± s.e.m of n = 3 biological replicates. Additional metabolites after shikimate feeding are shown in **Supplementary Figures 8 & 9**. Significant differences to Wt (R108) control are indicated; \**P* ≤ 0.05.

To determine if global metabolite changes occurred after shikimate feeding, additional amino acids and TCA pathway metabolites were analyzed by GC-MS. Interestingly, glutamine levels were higher in H_2_O-fed Wt, compared with mutants (**Supplementary Fig. 7**), and upon shikimate feeding, glutamine levels were significantly higher in Wt (**Supplementary Fig. 7**). The levels of TCA cycle intermediates may indicate the functionality of the Tyr catabolic pathway; however, those TCA cycle intermediates analyzed here, including fumararte, were not consistently altered after shikimate feeding in Wt and mutants (**Supplementary Fig. 8**). Citrate levels were reduced in *pdh1-1* and *pdh1-2* compared with Wt in H_2_O control treatment; however after shikimate feeding these differences were no longer apparent (**Supplementary Fig. 8**). These data did not provide evidence to support the involvement of the PDH enzyme in Tyr catabolism to the TCA cycle intermediate.

### PDH has limited role in the legume-rhizobia symbiosis

To test if PDH plays a role in legume-rhizobia symbiosis (**hypothesis IV**), Wt plants and *pdh1-1* mutants were grown side-by-side on low nitrogen Fahräeus medium and inoculated with a well-chracterized rhizobium of *M. truncatula*, *Sinorhizobium meliloti* Rm1021. Nodules were counted 14, 21, and 28 days post-inoculation (dpi) and acetylene reduction assays (ARA) were performed to measure nitrogenase activity (Wych and Rains, 1978). At all timepoints *pdh1-1* mutants did not display any phenotypic difference from Wt plants, including the number of nodules produced per root (**Fig. 5a**), suggesting that PDH is not essential for nodule development. *M. truncatula* forms indeterminate nodules that displayed the four standard zones: I, meristematic; II, infection; III, fixation; and IV, senescence zones in both Wt and *pdh1-1* (**Fig. 5b**, Van de Velde et al., 2006). Expression of the bacterial *nifH* gene, which is required for nitrogen fixation, was monitored through the use of a *S. meliloti* strain expressing a P*nifH::uidA* fusion to evaluate nodule maturation on *pdh1-1* and Wt plants (Starker et al., 2006). After 21 and 28 dpi, β-glucuronidase (GUS) activity and the overall nodule development were not altered between *pdh1-1* and Wt (**Fig. 5c**). Although nitrogenase activity in *pdh1-1* appeared to be slightly reduced compared with Wt at 21 and 28 dpi when nodule senescence might have been initiated, no significant differences was observed (**Fig. 5d**). ARA were repeated at the 21 dpi timepoint with *pdh1-1* and this time including *pdh1-2*. Nitrogenase activity was not significantly different between genotypes, even though *pdh1-1* showed a slight reduction, similar to the first experiment (**Fig. 5e**).

**Figure 5.**
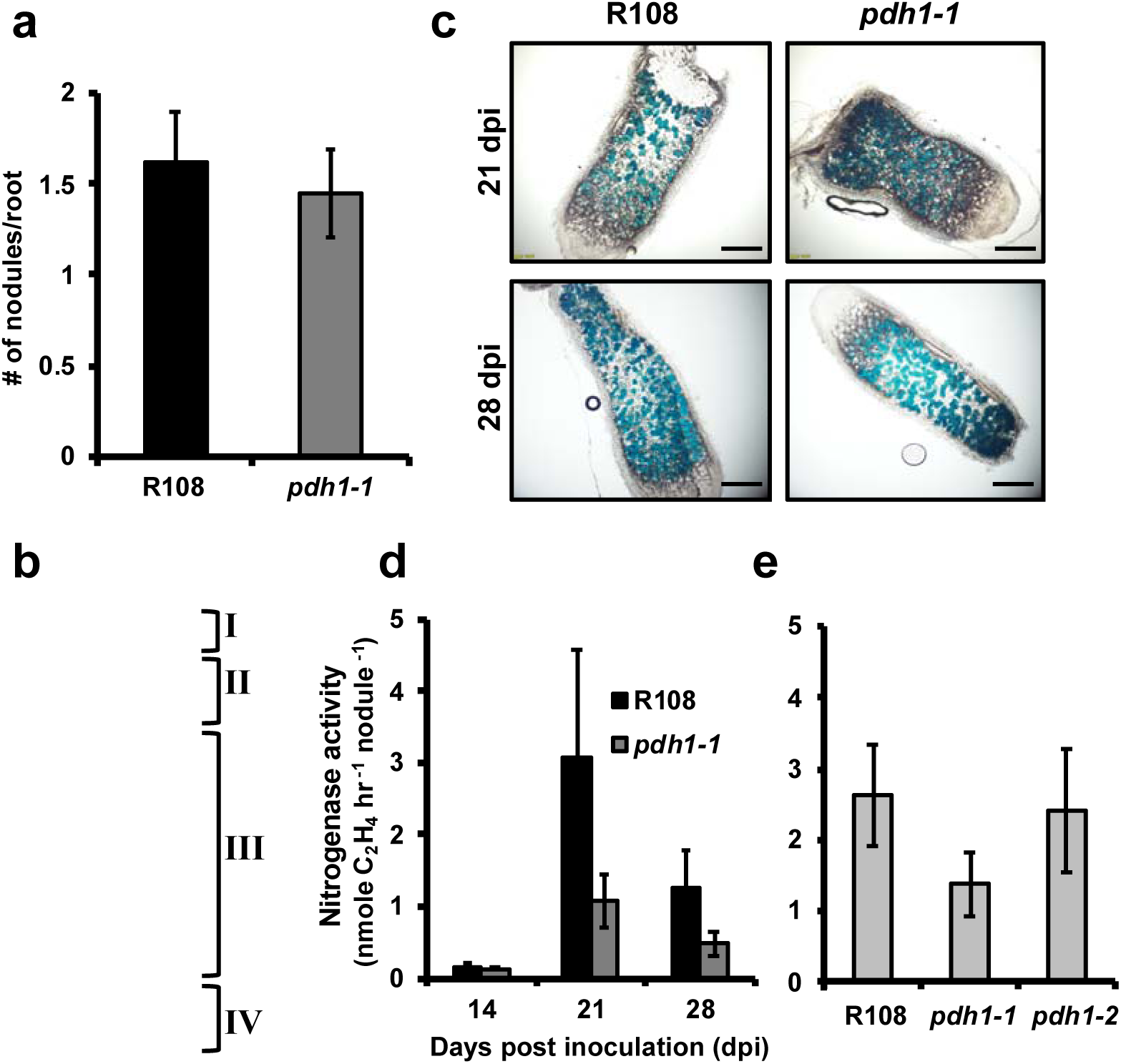
The role of PDH in legume-rhizobia symbiosis. **(a)** Wt and *pdh1-1* were grown on plates with low nitrogen Fahräeus medium, and 14 days post inoculation (dpi) with *S. meliloti* nodules were counted. Bars represent average number of nodules per root ± s.e.m. with n > 11 plants. **(b)** Indeterminate nodules from *M. truncatula* consist of four developmental zones. I meristem, II infection, III fixation, and IV senescence located closest to the root. These four developmental zones are shown on a characteristic nodule (enlarged for visualization) from 28 dpi. GUS staining with P*nifH*::UidA shows expression of bacterial *nifH* gene localized to the bacteroids (blue). **(c)** Thin sections of nodules from 21 and 28 dpi showing expression of P*nifH*::UidA, highlight bacteroid and nodule development in R108 and *pdh1-1*. Approximately equal staining was observed in all nodule developmental zones, suggesting *pdh1-1* is not affected in nodule development or bacteroid number, scale bar = 1 mM. **(d)** Nitrogen fixation efficiency was measured using an acetylene reduction assay (ARA). Plants were grown under the same conditions as in **a**, 14, 21 and 28 dpi ethylene production was measured and expressed as the average ± s.e.m with n > 7. **(e)** Nitrogen fixation efficiency assay performed in the same way as in **(d)** however, only at 21 dpi and with both mutant alleles and Wt. Activity is expressed as the average ± s.e.m with n > 4.

Independently, we also measured the presence and absence of PDH from plant tissue extracts of over twenty legume species, which were sampled across the legume phylogeny (Azani et al., 2017, **Supplementary Fig. 9**) and compared with their capacity to form nodules (Afkhami et al., 2018). All legumes sister to previously analyzed *G. max* (soybean) and *M. truncatula* (Schenck et al., 2015) showed PDH activity including *Arachis ipaensis* (peanut), which possess a TyrA enzyme with similar ADH and PDH activity (Schenck et al, 2017). All of these legumes were previously reported to be able to nodulate (Afkhami et al., 2018). When four legume species in the genistoid crown were analyzed (e.g. *Lupinus polyphyllus*), PDH activity was not detectable in any of them, despite successful detection ADH activity (**Supplementary Fig. 9**). Importantly, these legumes were reported to be able to nodulate, suggesting that legume plants having undetectable levels of PDH activity can still maintain nodulation. Some early diverging legume lineages are known to nodulate. Out of four of these species sampled from the mimosid crown three showed PDH activity (**Supplementary Fig. 9**). In contrast, all four species from caesalpinioid, another early diverging legume clade, are incapable of nodulating (Afkhami et al., 2018), but showed PDH activity (**Supplementary Fig. 9**). The lack of detectable PDH activity (e.g. in the genistoid crown) may be simply due to insuffient sensitivity with HPLC-based assay and/or the use of tissues that do not necessarily express *PDH*. However, detection of PDH activity in species lacking nodulation (e.g. in the caesalpinioid node) strongly support the lack of positive correlation between the presence of PDH activity and nodulation across the legume phylogeny. Phylogenetic sampling (**Supplementary Fig. 9**) together with *pdh1* mutant analysis (**Fig. 5**) together suggest that PDH is not essential for legume-rhizobia symbiosis, though more detailed analyses are needed to fully address its contribution to potentially enhance nitrogen fixation within the nodules in some legumes.

## DISCUSSION

In this study, we sought to understand the function of legume-specific PDH enzymes through analysis of *Tnt1*-transposon mutants of *PDH* in *M. truncatula*. Not only is *M. truncatula* a convenient model system having mutant populations and many genomic/transcriptomic resources, but *M. truncatula* also has a single *PDH* gene unlike some other legumes, i.e., soybean, having two *PDH* genes (Schenck et al., 2015). Analysis of two independent *pdh1* mutant lines suggests that MtPDH1 is responsible for all the PDH activity in *M. truncatula* (**Fig. 1**). This observation is consistent with chromatographic separation of PDH from ADH in soybean, which resulted in the detection of a single PDH peak as compared to multiple ADH peaks (Schenck et al., 2015). Our genetic data also support that ncADH, and ADH enzymes do not contribute to PDH activity, in agreement with *in vitro* data of soybean and *M. truncatula* enzymes (Schenck et al., 2015; Schenck et al., 2017a).

Surprisingly, the null *pdh1* mutants, displayed no visible aberrant growth phenotypes (**Fig. 1**). A slight bushy phenotype was observed in the *pdh1-2* allele, but not in *pdh1-1*, and thus here we focused on consistent responses observed in both alleles to avoid potential pleiotropic effects specific to *pdh1-2*. The lack of substantial phenotypes in *pdh1* is in contrast to the highly compromised growth and leaf development phenotypes of Arabidopsis *adh2/tyra2* mutant (de Oliveira et al., 2019). In *M. truncatula*, we were also unable to recover homozygous mutants of the canonical *ADH*, which is a single copy gene in *M. truncatula* as compared to two *ADH* copies present in Arabidopsis (Rippert and Matringe, 2002), further suggesting the essential nature of the ADH-mediated Tyr pathway in legumes. Thus, the canonical plastid-localized ADH pathway is the predominant Tyr biosynthetic route and cannot be compensated by the cytosolic PDH pathway in legumes.

Using the *pdh1* mutants, this study further systematically evaluated potential roles of PDH in Tyr biosynthesis (**hypothesis I**), biosynthesis of Tyr- and HPP-derived metabolites (**II**), Tyr catabolism and senescence (**III**), and legume-rhizobia symbiosis (**IV**). The obtained data provide evidence for support of **hypothesis I** that the PDH enzyme is involved in Tyr biosynthesis when carbon flux is increased through the shikimate pathway. Although Tyr levels were not altered between Wt and mutants under standard conditions (**Fig. 3a**), upon feeding with shikimate, Wt accumulated over 2-fold more Tyr than *pdh1-1* and *pdh1-2* (**Fig. 4**). Although AAA biosynthesis is localized to the plastids (Bickel et al., 1978; Jung et al., 1986; Maeda and Dudareva, 2012), some isoforms of the shikimate and AAA pathway enzymes were detected in the cytosol (Rubin and Jensen, 1985; Ganson et al., 1986; Ding et al., 2007). Cytosolic CM activities have been detected from various plant species including soybean (Goers and Jensen, 1984; Kuroki and Conn, 1989; Mobley et al., 1999; Schenck et al., 2015), which are not regulated by the AAAs, unlike their plastidic counterparts (Eberhard et al., 1996; Mobley et al., 1999; Westfall et al., 2014). Tyr-insensitive PDH enzymes provide a cytosolic route for conversion of prephenate into HPP, which can be further transaminated to Tyr by cytosolic Tyr-AT enzymes (**Fig. 1**) (Wang et al., 2016). Although the PDH-mediated Tyr pathway is not the predominant route for Tyr biosynthesis, it may provide an alternative route for additional Tyr synthesis under some conditions that increase flux through the shikimate pathway.

The current data also partly support **hypothesis II** that the PDH enzyme contributes to the production of specialized metabolites derived from Tyr and/or HPP in legumes. Although Tyr/HPP-derived tocopherols were not altered in the *pdh1* mutants under standard growth conditions (Fig. 3b), high light-induced tocopherol accumulation was partially attenuated in the *pdh1* mutant (**Supplementary Figure 3**). Therefore, the PDH enzyme may also contribute to tocopherol biosynthesis under stress conditions. Some specialized metabolic pathways emerged through duplication and neofunctionalization of genes from primary metabolic pathways (Weng et al., 2012; Moghe and Last, 2015). The legume PDH enzyme emerged as the result of a recent gene duplication of an *ADH* gene followed by a shift in substrate specificity from arogenate to prephenate (Schenck et al., 2017a). Thus, legumes may be able to divert the shikimate pathway flux to provide additional HPP or Tyr in the cytosol for the synthesis of downstream specialized metabolites including tocopherols. To further test this possibiliy, additional metabolomics experiments were conducted, but did not identify any putative HPP-derived compounds that were absent or lower in *pdh1* mutants than Wt (**Supplementary Table 1**). Although there are many Tyr and HPP-derived specialized metabolites produced in plants (e.g., rosmarinic acid, betalains, dhurrin, and benzylisoquinoline alkaloids), none appear to be specific to the legume lineage or correlate with the distribution of PDH activity (**Supplementary Fig. 9**) (Schenck and Maeda, 2018). Some species of the legume genus *Inga* can accumulate Tyr to 20% of leaf dry weight, which deters insect predation (Lokvam et al., 2006). Further analysis identified *Inga* species with high *PDH* expression that have hyper-accumulation of Tyr-derived specialized metabolites, such as Tyr-gallates (Coley et al., 2019). Therefore, more comprehensive analyses of Tyr-derived metabolites under different conditions, tissues types, and other legume species may identify specialized metabolites derived from the PDH enzyme in legumes.

The current data did not provide sufficient support for **hypothesis III** that the PDH pathway potentially contributes to Tyr catabolism and senescence. As Tyr is the most energetic amino acid (Hildebrandt et al., 2015), its catabolism is a crucial process in recovering energy. Mutations in Tyr catabolic genes can have dramatic effects of plant development (Han et al., 2013). Tyr is catabolized into fumarate, a TCA pathway intermediate, first by conversion into HPP via a Tyr-AT enzyme and further through the canonical Tyr catabolism pathway (**Supplementary Fig. 5a**) (Hildebrandt et al., 2015). Knockout of a soybean *HGO* did not cause lethality but led to an increase in vitamin E content by 2-fold, suggesting that a significant amount of carbon flux is diverted into vitamin E production when Tyr catabolism is blocked (Stacey et al., 2016). Furthermore, *MtPDH1* was co-expressed with *HPPD*, which together provide a direct pathway for homogentisate production. This potential coordination of PDH and HPPD, and cytosolic localization, can bypass three enzymatic steps of the ADH-mediated Tyr biosynthetic pathway, catalyzed by PPA-AT, ADH, and Tyr-AT to enter into the canonical Tyr catabolism pathway (**Fig. 1 & Supplementary Fig. 5a**). Additionally, *PDH* was co-expressed with many senescence-related genes (**Supplementary Fig. 1**, Van de Velde et al., 2006; Kusaba et al., 2013; Xi et al., 2013) suggesting that *PDH* may function when Tyr catabolism is enhanced. Despite these correlative data, however, experiments designed here to test the potential link between PDH and senescence did not provide evidence to support this hypothesis: upon dark-induced (**Supplementary Fig. 6**) and natural senescence (**Supplementary Fig. 4**) conditions that likely stimulate Tyr catabolism, no phenotypic differences were observed between mutants and Wt. Also, genes involved in Tyr catabolism were not consistently altered in mutants as compared to Wt (**Supplementary Fig. 5b**), and PDH expression and enzymatic activity were not induced upon senescence (**Supplementary Fig. 4c**). Thus, further experiments under different conditions (e.g. specific stress that induces Tyr catabolism) are needed to address the potential role of PDH in Tyr catabolism and senescence.

The data obtained in this study failed to directly support the **hypothesis IV** that the PDH pathway plays a role in legume-rhizobia symbiosis. Legumes initiate a symbiotic relationship with soil-dwelling rhizobia when there is insufficient nitrogen (Oldroyd et al., 2011). A chemical communication ensues that ultimately results in compatible rhizobia invading legume roots and formation of a new organ, the root nodule (Oldroyd, 2013). In the nodule, rhizobia fix atmospheric dinitrogen into ammonium, which is assimilated by the plant through the glutamine synthetase-glutamine oxoglutarate aminotransferase (GS-GOGAT) cycle, in which glutamine and glutamate are key amino acid carriers (Krapp, 2015). Mutations in *MtPDH1* did not alter nodule numbers (**Fig. 5a**) or the developmental progression of the nodules (**Fig. 5c**). Furthermore, nitrogenase activity was not significantly affected in mutants at any stage during the symbiotic interaction even when senescence might have been initiated (**Figs. 5d,e**). Alterations in nitrogen assimilation can lead to reduced nodulation in legumes (Streeter and Wong, 1988; Matamoros et al., 1999): for example, plants treated with a GS inhibitor phosphinothricin (PPT) result in loss of nodulation (Seabra et al., 2012) and also stimulate *MtPDH1* expression (**Supplementary Fig. 1**) as well as many other genes. Interestingly, *pdh1-1* and *pdh1-2* had reduced glutamine levels in the H_2_O-treated control leaves after 8 hours (**Supplementary Fig. 7**), which also persisted after shikimate feeding (**Supplementary Fig. 7**). Reduced glutamine levels in the *pdh1* mutants may, in turn, affect the symbiotic efficiency with rhizobia, although no statistically-significant reduction in nitrogenase activity was observed (**Figs. 5d,e**). Furthermore, PDH activity, which was measured in a phylogenetically diverse group of legumes, does not correlate with ability to nodulate (Azani et al., 2017; Afkhami et al., 2018, **Supplementary Fig. 9**). Thus, although PDH may provide an adaptive advantage to some legumes, it is likely not directly involved in and essential for the legume-rhizobia symbiosis.

## Acknowledgements

This work was supported by grants from the National Science Foundation IOS-1354971 to H.A.M. and NSF#1331098 and NSF#1546742 to JMA. LWS is supported in part by NSF awards 1340058, 1743594, 1139489, and 1639618. The University of Missouri, Office of Research provided initial instrumental and personnel funding for the MU Metabolomics Center. Development of *M. truncatula Tnt1* mutant population was, in part, funded by the National Science Foundation, USA (DBI-0703285 and IOS-1127155) to KSM. We thank the Germplasm Resources Information Network (GRIN) for providing some of the legume seeds used in this study.

## Conflict of Interest Statement

The authors declare no conflicts of interest.

## Author Contribution

CAS, JW, DJ, and KG carried our experiments and interpreted results. CAS, JMA, LWS, and HAM designed experiments. KSM and JW developed *Tnt1* mutant lines. CAS and HAM wrote the manuscript. All authors read and edited the manuscript.

## MATERIALS AND METHODS

### Plant Materials and Growth Conditions

*Medicago truncatula* seeds were scarified in concentrated hydrochloric acid for eight minutes and repeatedly washed with water. Seeds were then surface sterilized with bleach for 1.5 minutes followed by repeated water washes. Seeds were placed in sterile water for 16 hours at 4°C and then transferred to germination media (0.5x MS media, 0.8% agar, 1 μM GA_3_, pH 7.6), wrapped in aluminum foil and placed at 4°C. After 48 hours plates with sterilized seeds were moved to 22°C. After 24 hours the aluminum foil was removed and the germinated seedings were transferred to standard potting soil and placed in a growth chamber (Conviron). Pots were watered with 1x Hoaglands solution when dry, and grown under 12 hour light:dark cycles with 200 μE light intensity and ~60% humidity.

### Genotyping

A single young leaf from 6-week-old plants was placed in 1.7 mL microcentrifuge tube with 600 μL of DNA extraction buffer (10 mM Tris-HCl pH 8.0, 25 mM ethylenediaminetetraacetic acid (EDTA), and 0.5% SDS), pulverized with a mini blue pestle, and incubated at 55°C for 15 minutes. The solution was cooled to room temperature and 200 μL of 5 M ammonium acetate was added, vortexed for 20 seconds and centrifuged at 14,000 g for 3 minutes. The supernatant was transferred to a fresh tube and 600 μL of isopropanol was added, followed by centrifugation at 14,000 g for 1 minute. The supernatant was decanted and the pellet washed with 400 μL of 70% ethanol followed by centrifugation at 14,000 g for 1 minute. The supernatant was decanted and the pellet dried for 1 hour in a sterile hood. The resulting DNA was dissovled in 50 μL of H_2_0. Genotyping PCR reactions contained 1 μM gene and insertion specific primers (**Supplementary Table 2**), 1x EconoTaq PLUS master mix (Lucigen), and genomic DNA. DNA was amplified in a thermocycler with the following conditions: an initial denaturation at 95°C for 5 min, 35 cycles of amplification at 95°C for 20 s, 60–65°C for 20 s, 72°C for 60 s, with a final extension at 72°C for 5 min.

### Quantitative reverse transcription PCR (qRT-PCR)

RNA was extracted from leaves of 6-week-old plants. About 50mg of tissue was pulverized in liquid N_2_ using a mini pestle. RNA extraction buffer (68mM sodium citrate, 132mM citric acid, 1mM EDTA and 2% SDS) was added and vortexed immediately for 10 seconds, the tubes were then placed on their sides for 5 minutes at 22°C. The solution was centrifuged at 12,000g for 2 minutes and 400 μL of the supernatant was transferred to a fresh tube. 100 μL of of 1M NaCl was added and mixed by pipetting, followed by addition of 300 μL of chloroform, inverted multiple times and centrifuged at 4°C for 10 minutes at 12,000g. The upper phase was transferred to a new tube and an equal volume of isopropanol was added, mixed by inverting and placed at 4°C for 10 minutes. The solution was then centrifuged at 4°C for 10 minutes at 12,000g and the supernatant decanted. The pellet was washed with 70% ethanol, followed by centrifugation for 3 minutes at 12,000g and the supernatant decanted and the pellet dried in a sterile hood until all residual ethanol was evaporated. The resulting pellet was redissolved in 25 μL of nuclease free H_2_0 (Promega). To remove DNA, the RNA solution was treated with DNase (Turbo DNase, Fisher) following the manufacturer’s protocol. The remaining RNA was quantified using a nano-drop spectrophotometer (Thermo) and diluted to a 20 ng/μL concentration. RNA was converted into cDNA using reverse transcriptase (Applied biosystems) with an oligo d(T) primer.

For qPCR, cDNA was diluted to 5 ng/μL and additional 5-fold dilutions were made to calculate primer efficiency. All primer pair efficiencies were between 90-100%. cDNA was mixed with GoTaq qPCR master mix (Promega) containing SYBR green and 300 nM of each primer. Reactions were placed in an Stratagene Mx3000P (Agilent) thermocycler using the following PCR cycle an initial denaturation at 95°C for 10 min, 45 cycles of amplification at 95°C for 15 s, 60°C for 30 s, 72°C for 30 s.

For relative quantification, Ct values were extracted for each reaction and used to quantify initial cDNA concentration using 2^−ΔΔC(t)^ method normalized to a housekeeping gene (*MtPI4K* Kryvoruchko et al., 2016, **Supplementary Table 2**) using the LinRegPCR program (Ruijter et al., 2013). For absolute quantification, pET28a vectors carrying *MtPDH1*, *MtncADH* and *MtADH* were used as the template to amplify a fragment of the corresponding genes. The resulting fragments were purified from 0.8% agarose gels using a QIAquick gel extraction kit (Qiagen) following manufacturer’s protocol. The DNA concentration was quantified and repeated 5-fold dilutions were made and used to obtain a standard curve in qRT-PCR with gene-specific primers (**Supplementary Table 2**). Ct values were extracted for each qRT-PCR reaction and used to quantify initial cDNA concentration by using the linear range created as above for the respective gene.

### Enzyme extraction and ADH and PDH assays

Leaf tissue from 6-week-old plants were ground to a fine powder in a prechilled mortar and pestle under liquid N_2_. Extraction buffer (25 mM HEPES, pH 7, 50 mM KCl, 10% ethylene glycol, 1% polyvinylpyrrolidone (PVP) and 1 mM dithiothreitol (DTT)) was added in a 1:3 ratio of tissue to buffer (w/v). The slurry was centrifuged for 20 min at 4°C, at 20,000 g and the resulting supernatant desalted using a gel filtration column (Sephadex G50-80 resin, Sigma-Aldrich) equilibrated with extraction buffer without DTT and PVP. Protein concentrations were determined by a Bradford assay (Bio-Rad Protein Assay, Bio-Rad).

The desalted crude enzyme extracts were used in ADH and PDH reactions that contained 25 mM HEPES (pH 7.5), 50 mM KCl, 10% ethylene glycol, 1 mM NADP^+^, 1 mM substrate (L-arogenate or prephenate, respectively). Arogenate was prepared by enzymatic conversion from prephenate (Sigma-Aldrich), as previously reported (Maeda et al., 2011). Reactions were initiated by addition of enzyme from various sources and incubated at 37°C for 45 minutes. The resulting assays were injected into HPLC equipped with a ZORBAX SB-C18 column (Agilent) to directly detect the final product of the assay as described previously (Schenck et al., 2015).

For PDH and ADH activities from various legumes (**Supplementary Fig. 9**), leaf material was obtained from the University of Wisconsin-Madison Botany Department greenhouse or from identified trees on campus. Plants were grown under varying light and temperature conditions and leaf material was collected at different developmental stages. Enzyme extractions and ADH and PDH assays were performed as described above.

### Metabolite extraction and detection

Tissue from 6-week-old plants was added into 1.7 mL microcentrifuge tubes with 400 μL of extraction buffer (2:1 (v/v) methanol:chloroform, 0.01 % butylated hydroxytoluene (BHT), 100 μM norvaline and 1.25 μg/mL tocol as previously described (Collakova and DellaPenna, 2003), with 3 glass beads (3 mm). Samples were vigorously shaken for 3 minutes at 1000 r.p.m. using a genogrinder (MiniG 1600, SPEX SamplePrep). Additional chloroform (125 μL) and water (300 μL) were added and vortexed for 30 seconds. Samples were centrifuged for 10 minutes at 20,000 rpm and the polar and non-polar phases were transferred to new tubes and dried using a vacuum concentrator (Labconco).

For tocopherol analysis, the dried nonpolar phase was resuspended in methanol with 0.01% BHT. Samples were injected into a HPLC (Agilent 1260) equipped with a ZORBAX SB-C18 column (Agilent) using a 30 minute isocratic elution of 95 % methanol, 5 % water. Tocopherols in the extractions were visualized using fluorescence detection excitation at 290 nm and emission at 330 nm and compared with authentic standards (Sigma) and normalized to an internal control (tocol).

For anthocyanin detection, 1 M HCl of methanol was added to the polar phase in a 1:1 ratio. A spectrophotometer was used to measure absorbance at 520 nm. Absolute levels were estimated using an extinction coefficient of anthocyanin absorbance (Lee et al., 2005) of 33,000 L x M^−1^ x cm^−1^.

The polar and non-polar phases were analyzed with GC-MS. The dried polar phase was redissolved in pyridine with 15.0 mg/mL methoxyamine-HCl. Samples were vortexed for 30 seconds followed by sonication for 10 minutes and incubated for 60 minutes at 60°C. This was repeated once more, then the samples were derivatized with an equal volume of *N*-methyl-*N*-(tert-butyldimethylsilyl)trifluoroacetamide + 0.1 % *tert*-butyldimethylchlorosilane (MTBSTFA + 0.01 % t-BDMCS, Sigma-Aldrich) and incubated for 60 minutes at 60°C. Samples were then injected into GC-MS (Trace 1310, ISQ LT, Thermo Scientific). The dried non-polar phase was redissolved in 800 μL of chloroform with 50 ppm BHT and 500 μL of 1.25M HCl in methanol. Following incubation at 50°C for 4 hours, samples were completely dried under nitrogen gas. The dried non-polar phase was resuspended in 70 μL of pyridine and derivatized with 30 μL of MTBSTFA + 0.01 % t-BDMCS (Sigma-Aldrich). Following transfer to a glass vial, 1 μL of the polar and non-polar phases were injected onto a 30 m column (TG-5MS, Thermo Scientific) using a 10:1 split ratio, and an oven ramp method of 5°C per minute for 46 minutes and held at 300°C for 10 minutes. Detected compounds were compared with library matches from NIST and peak areas based on ion abundance were normalized to an internal standard (norvaline).

### Acetylene reduction assay (ARA)

Plants used for ARA were scarified and germinated as described previously. Seedlings were transferred to 12” square plates containing modified solid Fahräeus medium supplemented with 0.5 mM NH_4_NO_3_ (Catoira et al., 2000). Plates were grown under the same conditions as pot grown plants, however plates were placed at a ~60° angle so that the roots would not penetrate the agar. After 5 days of growth on plates, each plant was inoculated with 1 mL of water containing *Sinorhizobium meliloti* Rm1021 at an OD_600_ 0.02. Plants were then allowed to grow for 14, 21 and 28 dpi at which point nodules were counted and plants were moved to 10 mL glass jars with 1 mL of sterile water at the bottom. Glass jars were sealed with a rubber stopper, then injected with 1 mL of acetylene gas (10% acetylene final). After 48 hours of incubation at 37°C, 1 mL of the gas from the glass jars was injected into a gas chromatograph (GC-2010, Shimadzu) to measure the production of ethylene and and ARA activity was calculated as described in Hardy et al., 1968.

### Histochemical staining of lignin composition

For cell wall composition staining, thin cross sections of stem tissue from 6-week-old plants were prepared using a razor blade. For Mäule staining stem sections were placed in a 1% potassium permanganate solution for 5 minutes, then rinsed with water followed by addition of a 12% HCl (V/V) for 5 minutes and rinsed again with water. A 1.5% solution of sodium bicarbonate was added to facilitate a color change to dark red and visualized using an epifluorescence microscope (Olympus BX60).

For phloroglucinol staining, similar stem cross sections were placed in a well plate with 1 mL of 10 % phloroglucinol (w/v) solution in 95 % ethanol with 500 μL of 10 N HCl and incubated for 5 minutes. Stem sections were transferred to a glass slide and washed with water and visualized using a epifluorescence microscope (Olympus BX60).

GUS staining was performed on nodules developed after inoculation with *S. meliloti* Rm1021 carrying a P*nifH*::*UidA* fusion, to localize expression of the bacterial *nifH* gene, which is required for nitrogen fixation. Nodules were embedded in 4% agarose and 50-100 μM sections were made with a vibratome® 1000 plus (Leica). Sections were immersed in a staining solution (2.5% 5-Bromo-4-chloro-3-indoxyl-beta-D-glucuronic (X-Gluc), 0.2M sodium phosphate buffer (pH 7), 0.1 M potassium ferricyanide, 0.1 M potassium ferrocyanide, 0.25 M Na_2_EDTA, and 10% Triton X-100) and vacuum infiltrated for 10 minutes, incubated in the dark at 37°C for 30 minutes, and rinsed with phosphate buffer. Sections were visualized using bright field microscopy.

## SUPPLEMENTARY RESULTS

**Supplementary Table 1.** Non-targeted metabolite analysis from leaf tissue of Wt and *mtpdh1-1*

| Metabolite | retention<br>time (min) | Ion<br>(m/z) | pdh1-1/<br>R108 | pdh1-2/R1<br>08 | T-test<br>pdh1-1:<br>R108 | T-test<br>pdh1-2:R<br>108 |
| --- | --- | --- | --- | --- | --- | --- |
| **Beta-Alanine** | 21.5809 | 248.1 | NS | 1.49638 | NS | 0.04629 |
| **beta-D-glucoside** | 32.1517 | 204.1 | 1.18350 | 2.20999 | 0.01116 | 0.00016 |
| **Citric Acid** | 30.5877 | 273.1 | NS | 1.51560 | NS | 0.00027 |
| **D-(+)-Melibiose** | 43.5201 | 204.1 | 0.72128 | NS | 0.03323 | NS |
| **Ferulic Acid** | 36.1830 | 338.2 | NS | 0.51230 | NS | 0.01121 |
| **Gentisic Acid** | 29.4116 | 355.2 | NS | 2.85714 | NS | 0.00001 |
| **Glucuronic Acid** | 29.1234 | 292.2 | NS | 1.58440 | NS | 0.00001 |
| **Glycolic Acid** | 11.8422 | 205.1 | 1.42874 | 1.92904 | 0.01487 | 0.00037 |
| **L-Alanine** | 19.8168 | 188.1 | NS | 2.39806 | NS | 0.00005 |
| **Lauryl Alcohol** | 22.0027 | 243.1 | 1.27167 | NS | 0.01445 | NS |
| **Maleic Acid** | 18.2630 | 245.0 | NS | 0.68722 | NS | 0.04895 |
| **Malic Acid** | 23.0106 | 233.1 | NS | 0.73291 | NS | 0.04836 |
| **Malonic Acid** | 15.4680 | 233.1 | 0.75424 | NS | 0.02370 | NS |
| **Pinitol** | 30.8088 | 260.2 | 1.33875 | NS | 0.01147 | NS |
| **Propionic Acid** | 18.8585 | 292.2 | NS | 2.08458 | NS | 0.01253 |
| **Saccharic Acid** | 35.0341 | 333.2 | 0.74697 | 1.32846 | 0.00476 | 0.00815 |
| **Sucrose** | 44.9165 | 361.2 | NS | 1.99803 | NS | 0.02979 |
| **Tartaric Acid** | 25.6516 | 305.2 | 1.43056 | NS | 0.03744 | NS |
| **Trehalose** | 43.9602 | 361.2 | NS | 18.99309 | NS | 0.00048 |
| **Tyramine** | 32.7431 | 174.1 | NS | 0.59383 | NS | 0.00075 |
| **Unknown** | 33.6590 | 204.1 | 1.26433 | NS | 0.00363 | NS |
| **Unknown** | 17.5759 | 186.0 | 1.79713 | 2.62694 | 0.00452 | 0.01497 |
| **Unknown** | 13.6890 | 220.1 | 1.41853 | NS | 0.00884 | NS |
| **Unknown** | 30.8657 | 217.1 | 1.32711 | NS | 0.00953 | NS |
| **Unknown** | 9.2975 | 184.1 | 1.52453 | NS | 0.01137 | NS |
| **Unknown** | 47.8363 | 204.2 | 1.43690 | 1.35843 | 0.01504 | 0.04118 |
| **Unknown** | 47.2028 | 204.1 | 1.21975 | 1.20442 | 0.03006 | 0.04871 |
| **Unknown** | 32.5721 | 221.2 | 1.21109 | NS | 0.03112 | NS |
| **Unknown** | 49.3385 | 217.1 | 1.31458 | 1.40117 | 0.03151 | 0.01702 |
| **Unknown** | 36.3824 | 434.3 | 0.70531 | NS | 0.03570 | NS |
| **Unknown** | 21.0322 | 189.1 | 1.20140 | NS | 0.04070 | NS |
| **Unknown** | 23.6949 | 217.1 | NS | 3.31769 | NS | 0.00007 |
| **Unknown** | 32.7980 | 205.1 | NS | 3.21406 | NS | 0.00002 |
| **Unknown** | 27.4875 | 422.2 | NS | 3.08602 | NS | 0.01943 |
| **Unknown** | 17.5759 | 186.0 | NS | 2.48604 | NS | 0.01419 |
| **Unknown** | 24.6017 | 217.1 | NS | 2.38962 | NS | 0.00001 |
| **Unknown** | 22.0509 | 307.2 | NS | 2.33226 | NS | 0.00220 |
| **Unknown** | 19.1101 | 184.1 | NS | 2.02055 | NS | 0.04568 |
| **Unknown** | 30.6073 | 150.1 | NS | 1.50572 | NS | 0.00020 |
| **Unknown** | 24.9767 | 313.3 | NS | 1.43477 | NS | 0.04725 |
| **Unknown** | 31.7135 | 275.2 | NS | 1.31801 | NS | 0.00069 |
| **Unknown** | 38.4361 | 331.1 | NS | 1.27114 | NS | 0.00650 |
| **Unknown** | 33.7685 | 270.2 | NS | 0.76019 | NS | 0.03895 |
| **Unknown** | 41.0710 | 274.1 | NS | 0.70608 | NS | 0.01860 |
| **Unknown** | 37.2316 | 204.1 | NS | 0.67959 | NS | 0.02482 |
| **Unknown** | 25.7858 | 331.2 | NS | 0.65537 | NS | 0.01843 |
| **Unknown** | 31.0367 | 292.2 | NS | 0.54155 | NS | 0.00025 |
| **Xylose** | 26.8968 | 160.1 | NS | 0.60672 | NS | 0.00493 |

Only compounds are shown that were significantly different from Wt with abundance > 1.2 or <
0.8 fold-change and a P-value $\leq 0.05$ . Ratio of average relative metabolite abundance is shown
for *pdh1-1* and *pdh1-2* compared with Wt with N = 5. NS, not significant. Unknowns are
compounds that were detected with unique ions, but did not have a confident library match ( <

**Supplementary Table 2.**
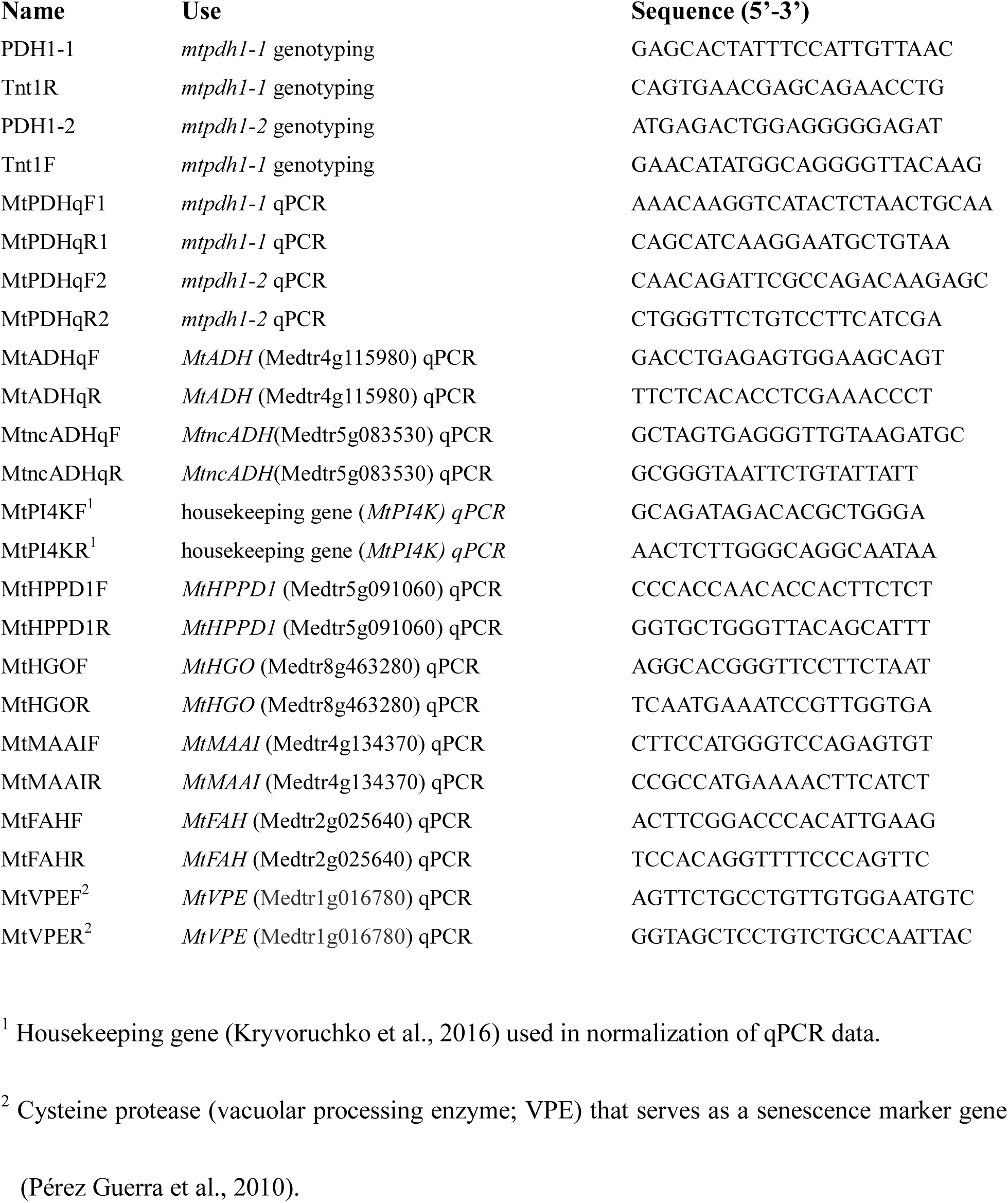
Primer sequences used in this study.

