## Supplementary Data for "Role of Cytosolic, Tyrosine-Insensitive Prephenate Dehydrogenase in *Medicago truncatula*"

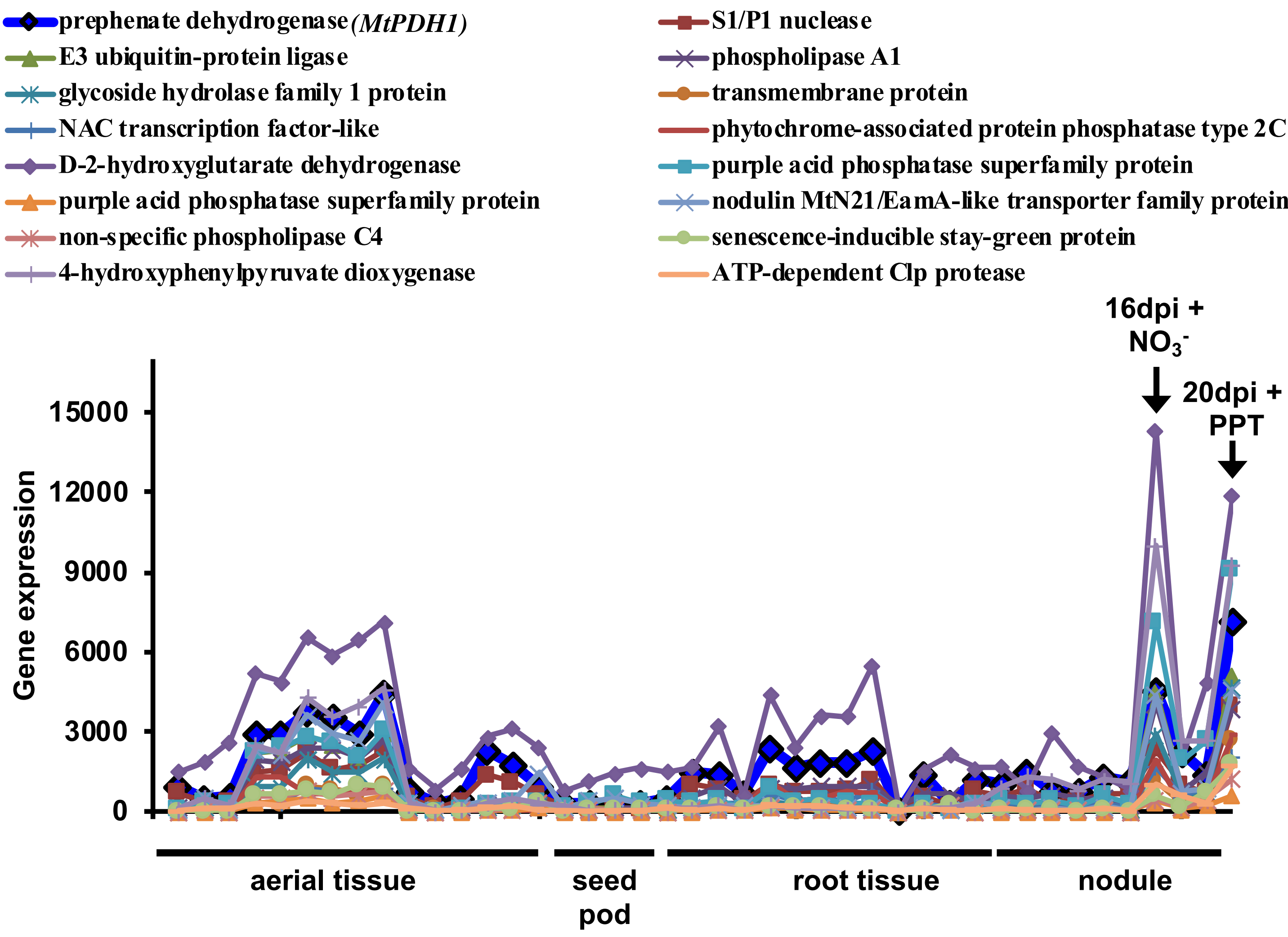


**Supplementary Figure 1. Co-expression analysis of *MtPDH*.** Co-expressed genes with *MtPDH1* were identified in various tissue-types and treatments using the Medicago gene expression atlas (www.mtgea.noble.org/v3/) with a Pearson's correlation coefficient of > 0.7. *MtPDH1* (blue line with square symbols) was co-expressed with senescence related genes, including proteases, lipases as well as HPP dioxygenase (HPPD), which catalyzes the initial step in Tyr catabolism. Additionally, *MtPDH1* was induced under two nodule conditions, highlighted by black arrows, that likely trigger senescence related processes, 16 dpi + nitrate ($NO_3^-$ Benedito et al., 2008) and 20 dpi + Phosphinothricin (PPT) treatment (Seabra et al., 2012).

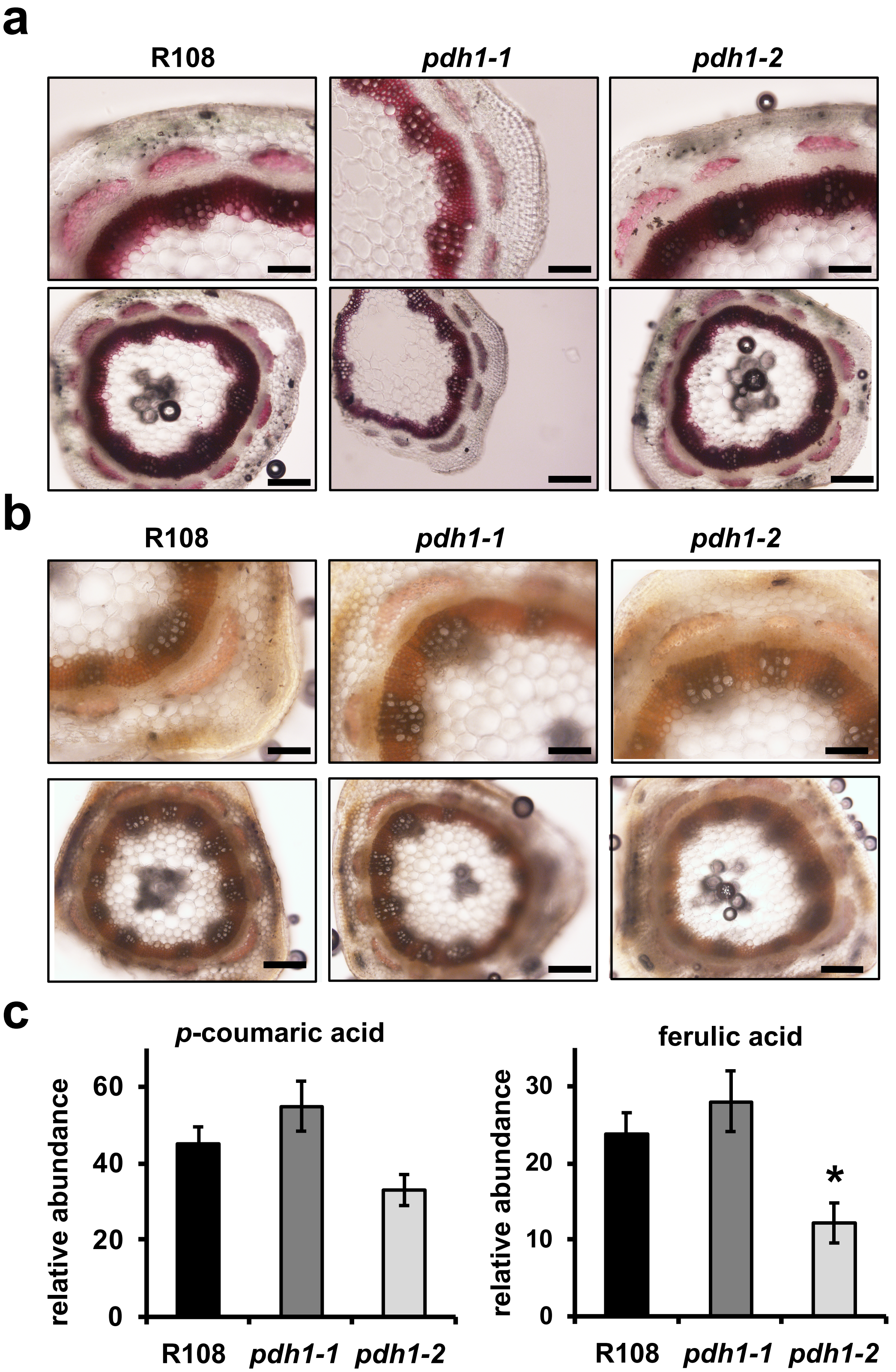


**Supplementary Figure 2. Effects of PDH pathway mutations on Phe-derived metabolism.** Lignin composition and linkages were assessed in Wt (R108) and mutants using two histochemical stains **(a)** phloroglucinol, and **(b)** Mäule staining. The top panel is 10x magnification (scale bars 50 μm), bottom panel is 4x magnification (scale bars 125 μm). Equal staining was observed in mutants compared with Wt (R108). **(c)** Phenylpropanoid compounds that serve as lignin precursors measured using GC-MS. Bars represent average metabolite abundance ± s.e.m of n ≥ 3 biological replicates. Significant differences to Wt control are indicated; $*P \leq 0.05$.

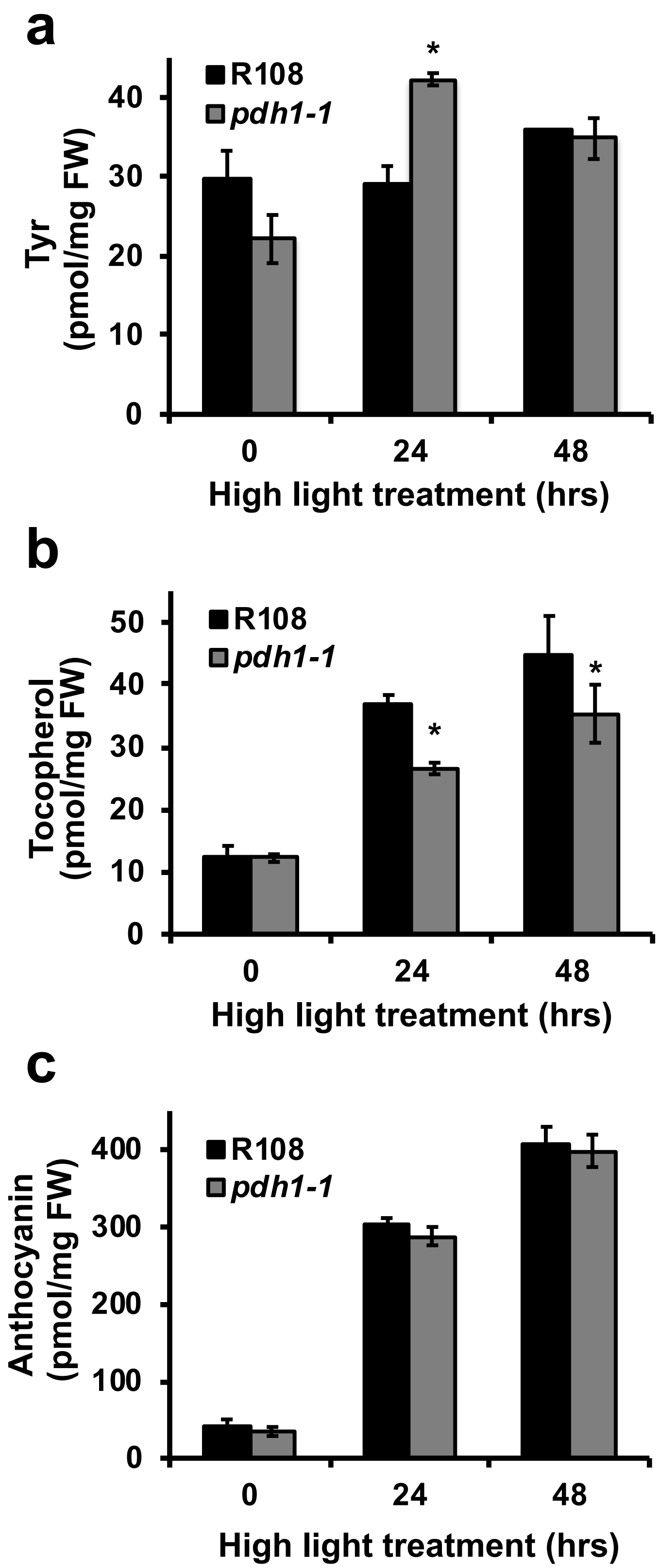


**Supplementary Figure 3. High light treatment stimulates Tyr- and Phe-derived metabolism.** Whole plants were moved from standard light (~200 μE) to high light (~950 μE) conditions and leaves from Wt (R108, black) and *pdh1-1* (gray) were collected after 0, 24, and 48hrs. Metabolites were extracted and identified using HPLC (Tyr and tocopherols) or spectrophotometrically (anthocyanins). Absolute quantification of Tyr **(a)**, tocopherols **(b)**, and anthocyanins **(c)** are shown as the average pmol/mg FW ± s.e.m of n = 3 biological replicates. Significant differences from the Wt (R108) control at the respective timepoints are indicated; $*P \leq 0.05$.

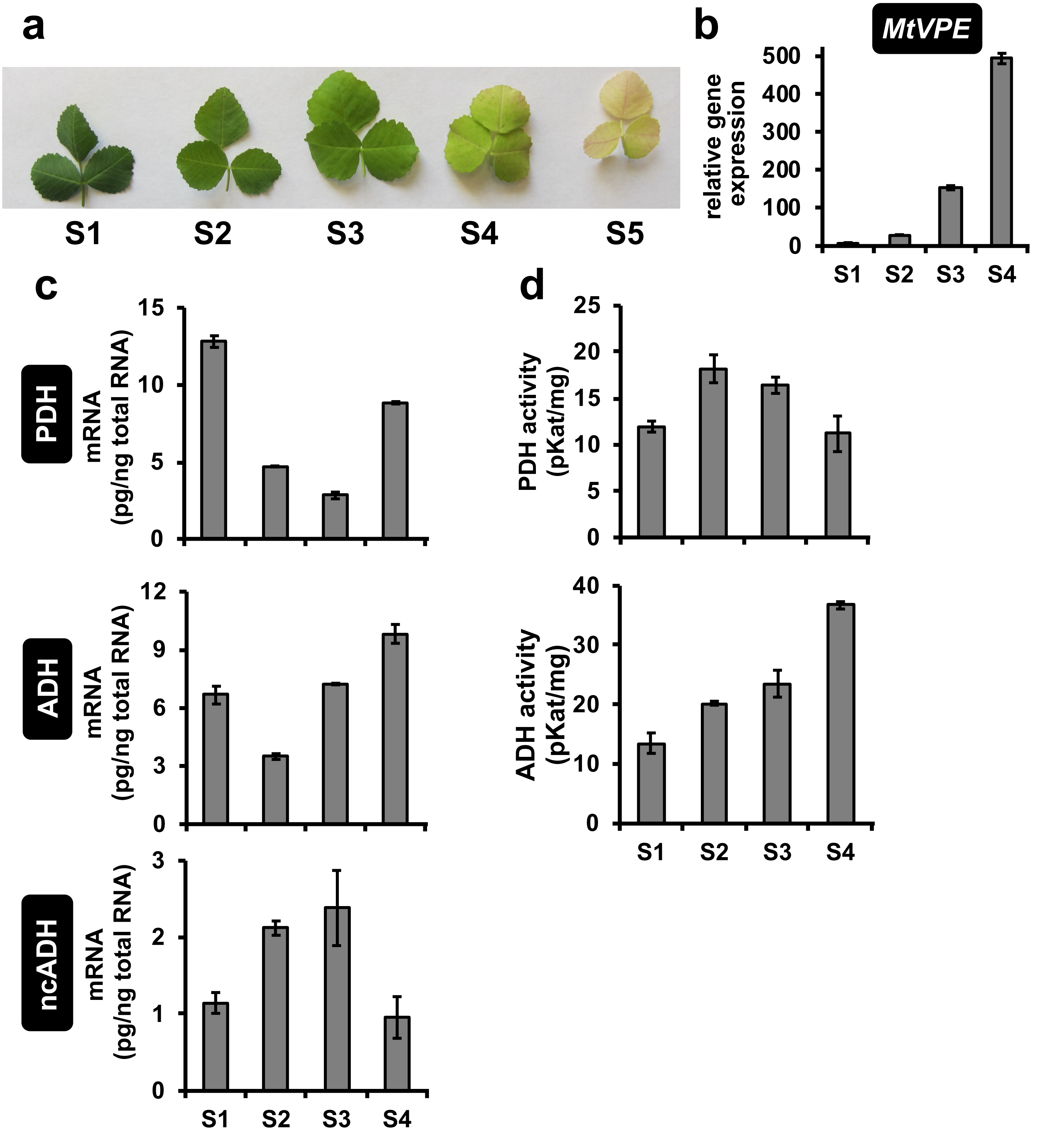


**Supplementary Figure 4. PDH and ADH gene expression and enzymatic activity in senescing leaves. (a)** Representative developmental time series of senescing Wt (R108) leaves used for gene expression **(b,c)** and enzymatic activity **(d)** analyses. High quality RNA and enzymes were obtained for all developmental stages except for S5. **(b)** A senescent marker gene (vacuolar processing enzyme, *MtVPE* Medtr1g016780*,* Pérez Guerra et al., 2010), was used as a control for qRT-PCR. Bars are average mRNA abundance ± s.e.m. of n = 3 biological replicates. **(c)** qRT-PCR analysis of *PDH* and *ADH* expression in senescing leaves, bars represent absolute mRNA abundance (pg/ng total RNA) ± s.e.m. of n = 3 biological replicates. **(d)** PDH and ADH enzymatic activity from senescing leaves. Bars represent average enzymatic activity (pKat/mg) ± s.e.m. of n = 3 biological replicates.

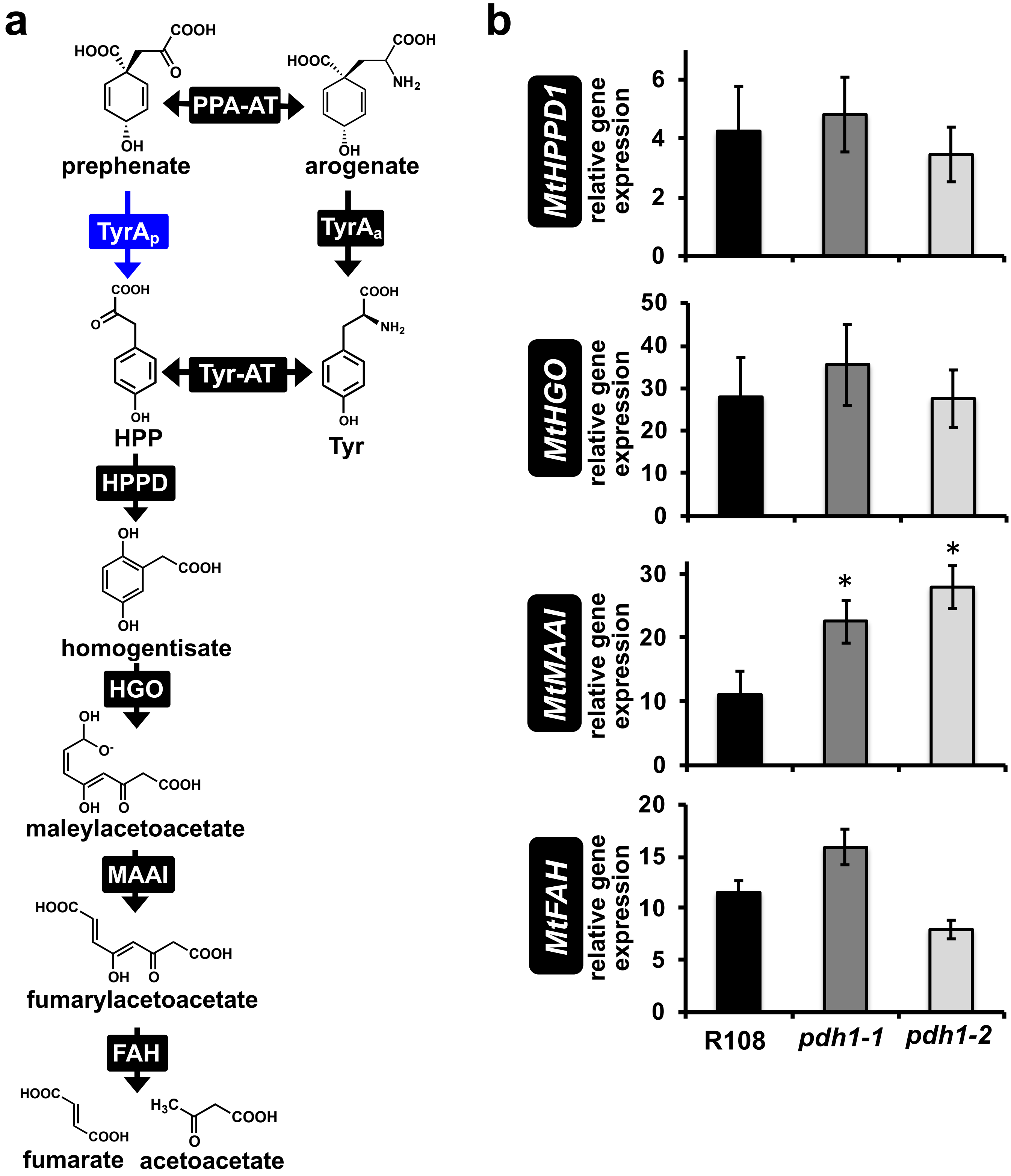


**Supplementary Figure 5. Tyr degradation pathway and gene expression in PDH mutants. (a)** The canonical Tyr catabolism pathway in plants. PDH (blue) provides a direct route to HPP and the Tyr catabolic pathway avoiding three ADH pathway steps (PPA-AT, $TyrA_a$, and Tyr-AT). **(b)** Gene expression analysis of Tyr catabolic pathway enzymes from RNA extracted from six week old leaf tissue from Wt (R108) and *mtpdh1* mutants. Bars represent average gene expression normalized to a housekeeping gene (*MtPI4K)* ± s.e.m. of n = 3 biological replicates. Significant differences to Wt (R108) control are indicated; **P* ≤ 0.05. FAH, fumarylacetoacetate hydrolase; HGO, homogentisate 1,2-dioxygenase; HPPD, HPP dioxygenase; MAAI, maleylacetoacetate isomerase; $TyrA_a$, arogenate dehydrogenase; $TyrA_p$, prephenate dehydrogenase; Tyr-AT; Tyr-aminotransferase.

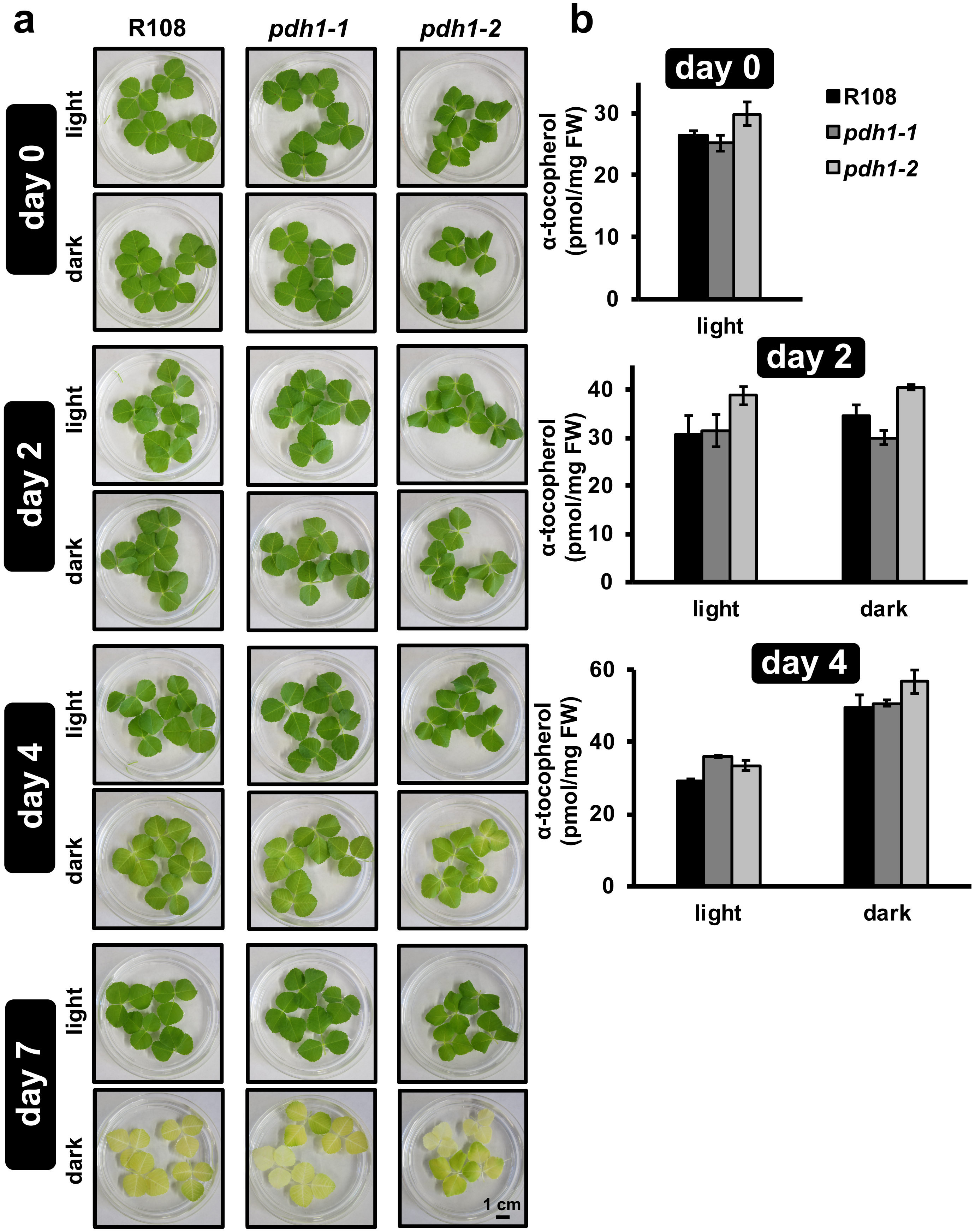


**Supplementary Figure 6. Dark-induced senescence leads to accumulation of tocopherols.** (**a**) Excised leaves from six week old Wt and *mtpdh1* mutants were floated on $H_20$ and treated in the dark, or left under normal light conditions for seven days. Dark-induced senescence progressed equally for Wt (R108) and mutants. (**b**) α-tocopherol content measured from leaves during the dark treatment expressed in pmol/mg FW ± s.e.m. of n = 3 biological replicates.

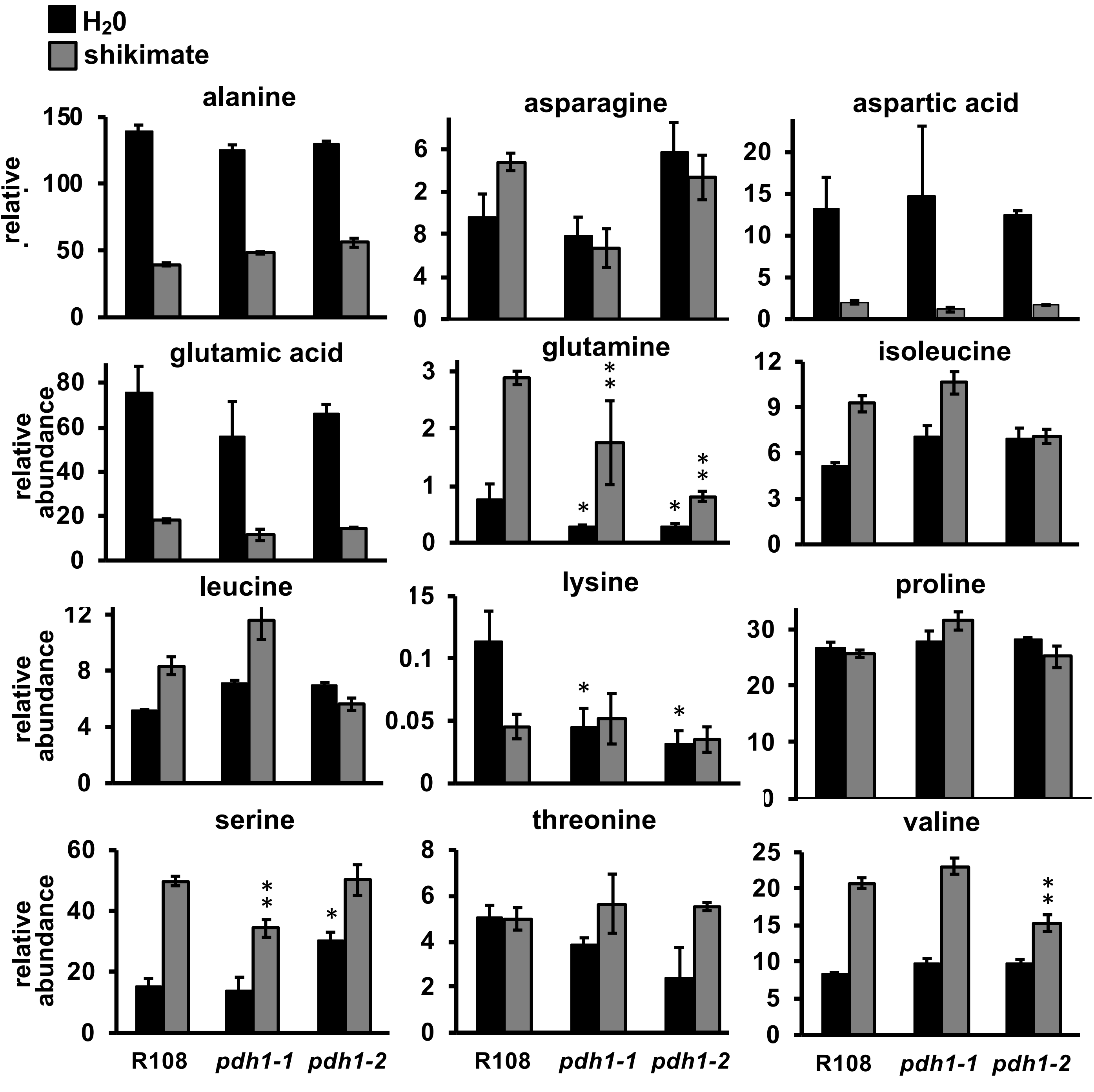


**Supplementary Figure 7. Additional amino acids from shikimate feeding in Wt and mutants.** Excised leaves from six week old plants were floated on $H_20$ (black bars) or a solution of 25 mM shikimate (Tyr precursor) (gray bars) for eight hours. Leaves were then used for metabolite extraction and identification using GC-MS. Relative abundance of the corresponding metabolites are the average ± s.e.m of n = 3 biological replicates. Significant differences from the water fed control (R108, black) are indicated; *$P \leq 0.05$, significant differences from the shikimate fed control (R108 gray) are indicated **$P \leq 0.05$.

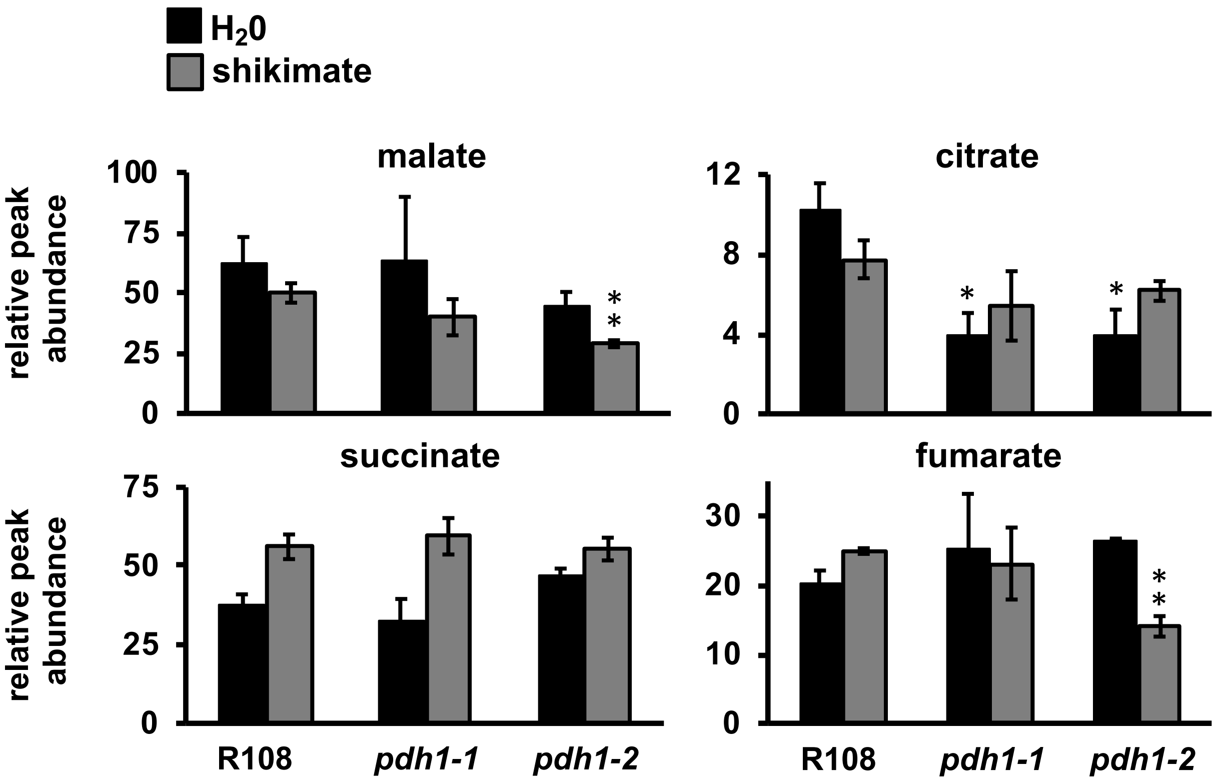


**Supplementary Figure 8. TCA metabolite levels following shikimate feeding in Wt and mutants.** Excised leaves from six week old plants were floated on $H_20$ (black bars) or a solution of 25 mM shikimate (Tyr precursor) (gray bars) for eight hours. Leaves were then used for metabolite extraction and identification using GC-MS. Relative abundance the corresponding metabolites are the average ± s.e.m of n = 3 biological replicates. Significant differences from the water fed control (R108, black) are indicated; $*P \leq 0.05$, significant differences from the shikimate fed control (R108 gray) are indicated $**P \leq 0.05$.

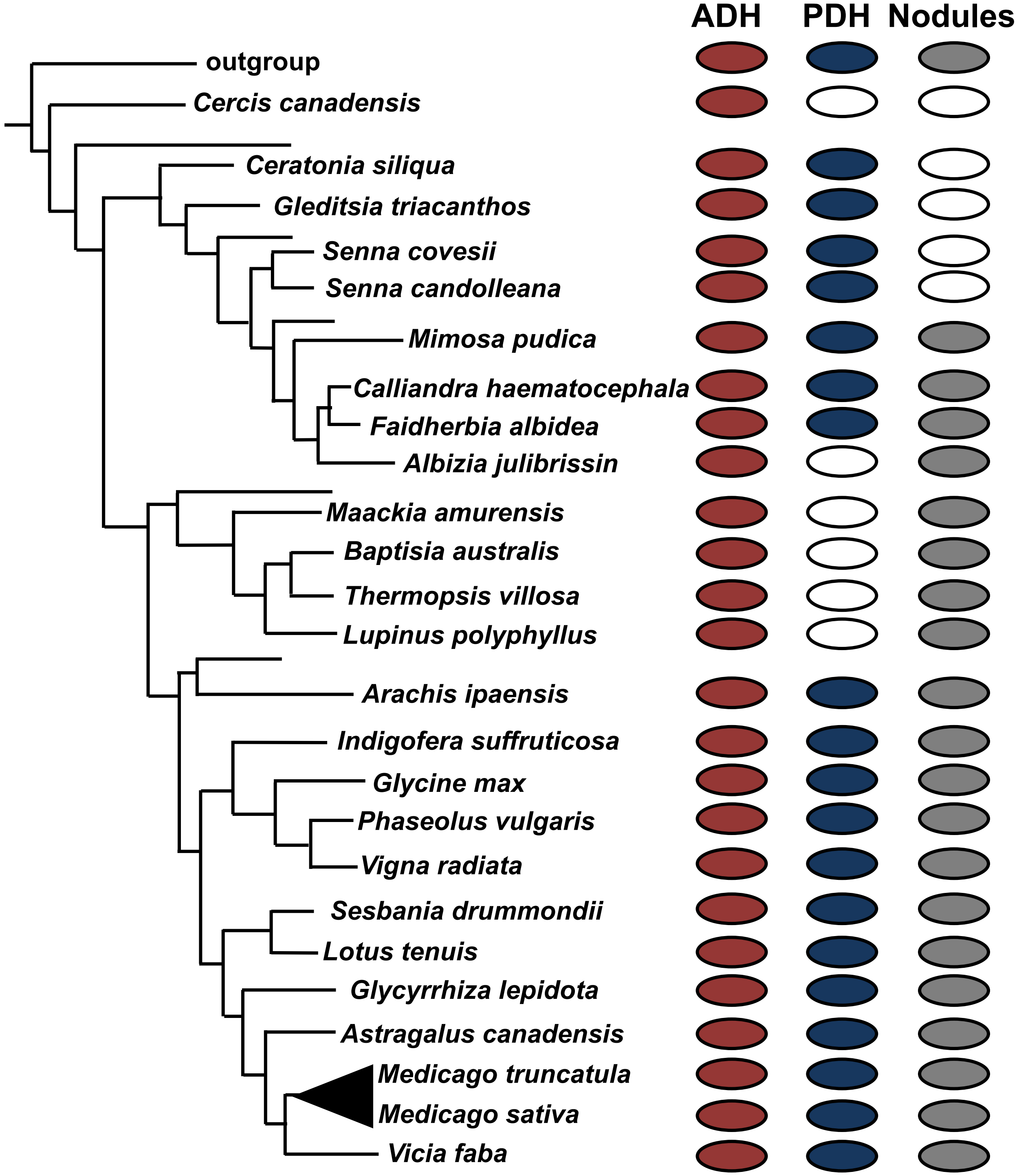


**Supplementary Figure 9. Phylogenetic distribution of ADH and PDH activity and nodulation status in Leguminosae.** ADH and PDH activities were assayed from various legumes and mapped onto a representative phylogeny of Leguminosae (Azani et al., 2017). Filled red circles indicate presence of ADH activity, filled blue circles indicate presence of PDH activity, whereas empty circles represent absence of enzymatic activity. Nodulation status was determined from (Afkhami et al., 2018). A filled grey circle indicates legumes that form nodules and an empty circle represents legumes that do not form nodules.
